# Histologic variants in bladder cancer harbor aggressive molecular features including TM4SF1 expression and a CA125+ cell state

**DOI:** 10.1101/2024.05.20.595039

**Authors:** Heiko Yang, Hanbing Song, Elizabeth Yip, Kevin Chang, Paul Allegakoen, Kevin L. Lu, Keliana Hui, Julia Pham, Corynn Kasap, Vipul Kumar, Janae Gayle, Bradley A. Stohr, Chien-Kuang Cornelia Ding, Arun P. Wiita, Maxwell V. Meng, Jonathan Chou, Sima P. Porten, Franklin W. Huang

**Affiliations:** Department of Urology, University of California San Francisco, San Francisco, CA; Department of Medicine, Division of Hematology/Oncology, University of California San Francisco, San Francisco, CA; Department of Pathology, University of California San Francisco, San Francisco, CA; College of Letters and Science, University of California Santa Barbara, Santa Barbara, CA; Chan Zuckerberg Biohub, San Francisco, CA; Bakar Computational Health Sciences Institute, San Francisco, CA; Institute of Human Genetics, San Francisco, CA; San Francisco Veterans Affairs Medical Center, San Francisco, CA

## Abstract

Histologic variant (HV) subtypes of bladder cancer are clinically aggressive tumors that are more resistant to standard therapy compared to conventional urothelial carcinoma (UC). Little is known about the transcriptional programs that account for the morphological and biological differences in HV tumors. To investigate the tumor biology of HV bladder cancers, we generated a single cell RNA sequencing (scRNA-seq) atlas of nine HV tumors and three UC tumors. Our analyses revealed a tumor cell state specific to HVs that is characterized by expression of *MUC16* (CA125)*, KRT24*, and *WISP2*. This CA125+ cell state bears transcriptional hallmarks of epithelial-mesenchymal transition, is enriched in metastases, is predicted to be highly chemotherapy resistant, and is linked with poor survival, suggesting that this cell state plays an important role in the aggressive biology of HV tumors. Our analyses also provide novel evidence of transcriptional “mimicry” between HVs and histologically similar non-urothelial cell types. Lastly, we identified higher expression of TM4SF1, a cell surface protein associated with cancer metastasis, in HV tumor cells compared to UC tumor cells. Finally, CAR T cells engineered against TM4SF1 protein demonstrated *in vitro* and *in vivo* activity against bladder cancer cell lines in a *TM4SF1* expression-dependent manner, highlighting its potential as a therapeutic target in bladder cancer.

**One sentence summary:** Single cell RNA sequencing of primary bladder cancers identified a CA125+ cell state specific to histologic variants that is associated with aggressive biological features and TM4SF1 as a novel therapeutic target for histologic variant subtypes of bladder cancer which can be targeted by anti- TM4SF1 CAR T cells.

## INTRODUCTION

Histologic variant (HV) subtypes of bladder cancer are found in up to 25% of all bladder tumors. Compared to bladder tumors with pure urothelial carcinoma (UC) histology, tumors with HVs are associated with worse clinical outcomes.^1,2^ The optimal clinical management of HV bladder cancers remains challenging as many HV subtypes do not respond well to systemic therapy and treatment options are limited and therefore represents a major unmet need.^3–7^

While significant progress has been made to define the molecular characteristics of pure UC,^8–10^ much less is known about the biology of HVs. Fundamentally, it remains unclear whether each HV subtype should be regarded as a distinct entity or whether HVs share common features as a group. Some genomic alterations, such as *TERT* promoter mutations in micropapillary, plasmacytoid, and adenocarcinoma variants, appear to be more associated with HVs than UCs, while others, such as *CDH1* truncations in plasmacytoid variants, are thought to be subtype defining.^11–15^ Emerging evidence suggests that HV biology may not be governed solely at the genomic level, but transcriptional analyses based on bulk RNA sequencing data remain limited to date.^16,17^ This use of bulk RNA sequencing is not well suited to study HVs because it requires large sample sizes that are difficult to achieve in HVs, especially when considering heterogeneity related to number of individual subtypes.

To more deeply understand the aggressive biology of HVs and to identify potentially targetable molecular features, we hypothesized that single cell RNA sequencing (scRNA-seq) could distinguish the transcriptional profiles of HV-containing bladder tumors given their heterogeneity. Our results highlight the potential for scRNA-seq to advance precision cancer medicine approaches in rare, understudied tumors.

## METHODS

### Sample collection

We obtained a total of 15 fresh bladder tumor samples from patients undergoing surgery at our institution under IRB 10-04057. In patients undergoing transurethral resection of bladder tumor (TURBT), specimens were obtained using cold biopsy forceps. In patients undergoing radical cystectomy, specimens were obtained immediately upon removal of the bladder to minimize the effects of ischemia. Visible tumor was excised from the specimen after the bladder was opened according to standard pathology protocol. Clinical and pathological data are shown in Supplemental Table 1. All tissue was immediately placed in RPMI 1640 media on ice.

### Tissue dissociation

Mechanical tissue dissociation was performed using scissors and enzymatic dissociation was performed using 1000 U/mL Type IV collagenase (Worthington, Cat: LS004188) at 37 °C for 30 minutes. A single cell suspension was isolated using a 40 µm strainer, pelleted at 300 g, and reconstituted in RPMI 1640 media with 10% FBS. Viability and concentration were determined using acridine orange / propidium iodide on a LUNA automated cell counter (Logos Biosystems). The suspension was then adjusted for a target loading concentration of ∼50,000-100,000 live cells/mL.

### Single-cell RNA sequencing

cDNA library preparation was performed using the Seq-Well platform as previously described.^18,19^ Briefly, 10,000-20,000 cells were loaded onto a Seq-Well array containing 110,000 barcoded mRNA capture beads (ChemGenes, Ct: MACOSKO-2011-10(V+)). Arrays were sealed using a polycarbonate membrane (Sterlitech, Cat: PCT00162X22100) at 37 °C for 40 minutes. Cells were then lysed in lysis buffer (5 M guanidine thiocyanate, 1 mM EDTA, 0.5% sarkosyl, 1% BME) for 20 minutes at room temperature. Hybridization of mRNA to the beads was performed in hybridization buffer (2 M NaCL, 4% PEG8000) for 40 minutes. The beads were then collected and washed with 2 M NaCl, 3 mM MgCl_2_, 20 mM Tris-HCl pH 8.0, 4% PEG8000.

Reverse transcription was then performed using Maxima H Minus Reverse Transcriptase (ThermoFisher, Cat: EP0753) in Maxima RT buffer, PEG8000, template switch Oligo dNTPs (NEB, Cat: No447L), and RNase inhibitor (Life Technologies, Cat: AM2696) at room temperature for 15 minutes and then 52 °C overnight. Second strand synthesis was performed using Klenow Exo- (NEB, Cat: M0212L) in Maxima RT buffer, PEG8000, dNTPs, and dN-SMRT oligo for 1 hr at 37 °C. Whole transcriptome amplification was performed with KAPA HiFi Hotstart Readymix PCR kit (Kapa Biosystems, Cat: KK2602) and SMART PCR Primer (Supplementary Data). The reactions were purified using SPRI beads (Beckman Coulter) at 0.6X and then 0.8X volumetric ratio.

Libraries were prepared using 800-1000 pg of DNA and the Nextera DNA Library Preparation Kit. Dual- indexing was performed using N700 and N500 oligonucleotides. Library products were purified using SPRI beads at 0.6X and then 1X volumetric ratio. A final 3 nM dilution was prepared for each library and sequenced on an Illumina NovaSeq S4 flow cell.

### Sequencing and alignment

Sequencing results were returned as paired FASTQ reads. These paired FASTQ files were then aligned against the hg19 reference genome (GRCh37.p13) using the dropseq workflow (https://cumulus.readthedocs.io/en/latest/drop_seq.html). The alignment pipeline output for each pair of FASTQ files included an aligned and corrected bam files, a digital gene expression (DGE) matrix text which was used for downstream analysis, and text-file reports of basic sample qualities such as the number of beads used in the sequencing run, total number of reads, alignment logs.

### Single-cell quality control and clustering analysis

Cells were clustered and analyzed using Seurat (v4.3.0) in R (v.4.3.1). Cells with fewer than 300 genes, 500 transcripts, or a mitochondrial gene content of 20% or greater were removed. Doublets were removed using DoubletFinder (v.2.0.3). UMI-collapsed read-count matrices for each cell were used for clustering analysis in Seurat. We followed a standard workflow by using the “LogNormalize” method that normalized the gene expression for each cell by the total expression, multiplying by a scale factor 10,000. To identify different cell types, we computed the standard deviation for each gene and returned the top 2,000 most variably expressed genes among the cells before applying a linear scaling by shifting the expression of each gene in the dataset so that the mean expression across cells was 0 and the variance was 1. Principal components analysis (PCA) was run using the previously determined most variably expressed genes for linear dimensional reduction and the first 100 principal components (PCs) were stored, which accounted for 47.04% of the total variance. For graph-based clustering, the top 75 PCs and a resolution of 0.5 were selected, yielding 36 cell clusters. Differentially expressed genes (DEGs) in each cluster were identified using the FindAllMarker function within the Seurat package and a corresponding p-value was given by the Wilcoxon’s Rank Sum test followed by an FDR correction. In the downstream analysis, tumor cells from each patient were further clustered in a similar manner. For the individual patient clustering analysis, the number of PCs was determined by the statistical permutation test and the straw plot, and clustering resolution was selected accordingly.

### Cell-type annotation and copy number variation

To annotate each cell type from the previous clustering, we referred to canonical markers and signature gene sets developed from established studies for each cell type. We computed the signature scores of these established gene sets for each cell in our dataset using the AddModuleScore function in Seurat. Each cluster in our dataset was assigned with an annotation of its cell type by top signature scores within the cluster. To validate the identities of the tumor cell populations, we estimated copy number variants (CNV) via InferCNV (Version 1.4.0), using all the non-tumor populations as reference. During the inferCNV run, genes expressed in fewer than five cells were filtered from the data set and the cut off was fixed at 0.1. Hidden Markov model (HMM) based CNV prediction was generated and estimated CNV events were shown in a heatmap.

### Pseudotime analysis

To further investigate the differential trajectories of tumor cells in each patient, we conducted a pseudotime analysis in Monocle3. To analyze gene expression relative to the Cluster 13 cell state, Cluster 13 cells were selected as the starting point for the pseudotime trajectory. Pseudotime trajectories were computed accordingly and visualizations were made to illustrate specific gene expression levels along the pseudotime trajectory in each patient.

### Gene ontology and gene set enrichment analysis

Within the tumor cells, we created a customized gene set signature for each variant tumor cell population of interest. Using the DEGs obtained from FindAllMarker function, we included genes with log2 fold change > 2 and statistical significance (FDR q < 0.05) in the customized signature gene set.

To assess the *in silico* functional roles of Cluster 13 cells, we used the signature gene sets derived from the scRNA-seq data to run gene ontology (GO) analysis against known signature gene set collections such as Hallmark, C2CP, C2CGP, C5GO and C6 oncogene collections (https://www.gsea-msigdb.org/gsea/msigdb/). The gene ratio and statistical significance levels from the overexpression test were calculated. Normalized gene expression data and variant tumor types as metadata were used in the GSEA analysis run on the GSEA software.^20,21^

To examine the association between signature gene sets or marker expression derived from our dataset and known basal/luminal signatures or canonical marker expression in the TCGA-BLCA bulk RNA sequeqncing dataset for validation, we performed ssGSEA (single set Gene Set Enrichment Analysis) by projecting the TCGA sample expression data onto the transcriptomic space defined by marker expression and established signature gene sets. For each target marker expression of target signature gene set, association was quantified via IC (information coefficient) and statistical significance was computed.

### Survival analysis

Within the TCGA-BLCA bulk RNA-seq dataset, the Cluster 13 signature score was computed on the normalized gene expression data for each sample. Samples were then divided into high and low groups based on the 20% percentile cutoff of the Cluster13 signature score. The overall survival (OS) distribution of both groups was compared by means of log-rank tests using the survfit function from the survival package (v3.3-1). The Kaplan-Meier (KM) survival curve was plotted using the survminer (v0.4.7) package.

### Histology and immunohistochemistry

FFPE bladder cancer tissue (from the scRNA-seq cohort and additional specimens) banked under IRB 10-04057 was sliced to 4 µm and mounted on positively charged Superfrost microscope slides. Hematoxylin and eosin (H&E) staining was performed using a standard method. CA125 immunohistochemistry (IHC) was performed using a clinically validated mouse monoclonal antibody (Signet, clone OC125) on an automated Ventana Benchmark Ultra IHC system using CC1 cell conditioning solution. TM4SF1 IHC was performed using a rabbit polyclonal antibody (Abcam, ab113504) at a 1:500 dilution after a 10-minute citrate antigen retrieval at 100 °C on a Leica Bond III platform. A tissue microarray including pancreas, vascular endothelium, adipose, and lymphoid tissue was used for positive and negative control. The signal in tumor cells was compared with that of endothelial cells on the same slide; tumor cells that stained equally or darker than endothelial cells were scored as “strong” while those that stained lighter were scored as “weak.” All TM4SF1 and CA125 stains were reviewed by a pathologist.

### CA125 serology

Serum CA125 levels were prospectively measured in patients undergoing TURBT or cystectomy for bladder tumors using the Abbott Architect Chemiluminescent Microparticle Immunoassay (CMIA) and reviewed under IRB 10-04057. Blood samples were drawn in the preoperative area prior to surgery. Pathologic diagnoses were reviewed. Tumors with >5% HV components were categorized as “HVs” while tumors with no mention of HV were categorized as “UC.” Tumors with equivocal or negligible HV components were excluded from the analysis; patients with “no tumor” on final pathology were also excluded.

### CAR constructs

The heavy (VH) and light (VL) chains of the TM4SF1 scFv binder was obtained from antibody AGX-A01 (patent US011208495B2). The VH and VL sequences were cloned in two configurations using the Gibson Assembly protocol (Twist) into a CAR backbone containing IgG4 spacers, CD8 hinge and transmembrane domain, 4-1BB costimulatory domain, CD3ζ chain, and EGFP. Plasmids were prepped using the NucleoBond Xtra Midi Plus kit (Takara Bio).

### CAR lentivirus production

For TM4SF1-CAR lentivirus production, HEK293T-Lenti-X cells (Takara Bio) were thawed, cultured, and expanded in DMEM media supplemented with 10% FBS. HEK293T-Lenti-X cells were transfected with the TM4SF1-CAR lentiviral plasmid and the packaging plasmids psPAX2 and pVSVG using the TransIT-LT1 transfection reagent (Mirus Bio). Cell supernatant was collected at 48 hours and 72h. The virus was filtered and concentrated using the Lenti-X Concentrator (Takara Bio) according to manufacturer’s instructions and resuspended in serum-free media.

### TM4SF1-CAR T generation

Human T cells were isolated from a leukopak (Stemcell Technologies) using the Easy Sep Human T cell enrichment kit (Stemcell technologies). T cells were then plated on retronectin coated plates (Takara, T100A), stimulated with Human CD3/CD28 T Cell Activator (Stemcell Technologies, 10971), and concentrated lentivirus was added. Cells with virus were spun at 1000 rpm for 45 minutes. After 72 hours of incubation, virus was removed, and cells were allowed to recover for 2-3 days. Transduction efficiency was evaluated via flow cytometry by GFP expression. If less than 30% of the T cells were GFP positive, the cells were MACs sorted using a biotinylated c-myc antibody (Miltenyi Biotec, 130-124-877) and isolated using the MiniMACS separator and columns (Miltenyi) according to manufacturer’s protocol. The CAR-T cells were grown in either ImmunoCult-XF T Cell Expansion Medium (Stemell Technologies, 01981) or TexMACS™ Medium (Miltenyi Biotech, 130-097-196). Human recombinant IL-15 (Stemcell Technologies, 78031) and IL-7 (Stemcell Technologies, 78053), 10 ng/mL final concentration each was freshly added to the cells every 2-3 days, with cells grown at a concentration of 1 × 10^6^ per mL and used between day 14 – 20 for downstream assays.

### Cell culture

5637 cells were obtained from the UCSF Cell Culture Facility. UMUC-3 cells were a gift from Bradley Stohr (UCSF). T24, UMUC-1 and 253JBV cells were gifts from Peter Black (University of British Columbia) and David McConkey (Pathology Core, Bladder Cancer SPORE, MD Anderson Cancer Center). Cells were grown in standard MEM media (Corning) supplemented with 10% FBS (Seradigm) and penicillin/streptomycin. All experiments were conducted within 20 passages from the parental stock. Cells were validated by STR profiling and routinely tested for mycoplasma (Lonza).

### TM4SF1 knockout cells

UMUC-3 TM4SF1-KO cells were generated by transient transfection (Lipofectamine 3000) of UMUC-3 cells with PX458 (Addgene, #48138). Each plasmid contained one of three different sgRNA targeting sequences: 1) AGTGCACTCGGACCATGTGG; 2) GGTGTAGTTCCACTGGCCGA; 3) ATTAGCCGCGATGCACAGGA. 48–72 h after transfection, GFP-positive cells were sorted by FACS (BD Fusion) and expanded. Cells were then stained with a TM4SF1 antibody (Miltenyi, clone REA851, 1:100), sorted a second time by FACS (BD Fusion), and negative cells were collected and expanded.

### TM4SF1 flow cytometry

Flow cytometric quantification of TM4SF1 expression across human bladder cancer cell lines was performed by incubating with anti-TM4SF1-PE antibody (Miltenyi, clone REA851, 1:100) for 30-60 minutes on ice. Cells were analyzed using an Attune NxT Flow and the median fluorescence intensity (MFI) was calculated and data were analyzed using FlowJo software.

### IncuCyte co-culture assays

Bladder cancer cells labeled with NucLightRed (Sartorius) were co-cultured with human non-transduced (NTD) T cells or TM4SF1-CAR T cells at variable effector-to-target (E:T) ratios. On day 0, 2000-5000 target cells were plated and allowed to adhere overnight. On day 1, effector T cells were added and tumor cell killing was monitored on an IncuCyte S3 (Sartorius). Images were obtained every 3-6 hours over 72-96 hours. Target cells were quantified based on the red object count or red area confluence normalized to the starting day 1 values, and data were plotted on Prism (GraphPad, v10).

### Animal studies

All animal studies were performed under an approved Institutional Animal and Use Committee (IACUC) protocol. NSG (NOD/SCID/gamma) mice were housed in the UCSF barrier facility under pathogen-free conditions and were obtained through an in-house breeding core. For subcutaneous xenografts, 1x 10^6^ cells were injected into the left flank of 8-10 week old male NSG mice. The injected cells were resuspended in 1:1 serum-free media and Matrigel (BD Biosciences). Mice were enrolled into treatment groups once tumor volumes reached between 50-100 mm^3^, typically 10-14 days after tumor cell inoculation. An intravenous injection of 3-5 x 10^6^ nontransduced (NTD) control or TM4SF1 CAR T cells was then delivered through the tail vein. Tumors were measured with digital calipers and mice were weighed twice weekly by personnel from the UCSF Preclinical Therapeutics Core in a blinded fashion. Tumor volumes were recorded using Studylog Animal Study Workflow software and plotted using Prism (GraphPad, v10). Mice were euthanized when tumors reached 20mm in any direction. For survival analysis, a log-rank test was used to compare the overall survival of mice in each cohort.

## RESULTS

### Single cell analysis of tumor epithelial cells reveals a novel CA125+ tumor cell state in histologic variants

We collected tissue and dissociated single cells from 4 pure urothelial carcinomas (UC) and 11 variant tumors. Detailed clinical information is displayed in Supplemental Table 1; pathologic diagnoses were confirmed in specimens collected for sequencing (Fig. S1). Single-cell RNA sequencing (scRNA-seq) was performed using the Seq-well platform, and the sequencing results were processed in our customized analytical pipeline (Fig. S2A). After ambient RNA decontamination and removal of low-quality cells, 21,533 cells in total were captured for downstream analysis from these specimens (Fig. S2B). While tumor epithelial cells were captured from almost all tumors, the capture rate for stromal and immune cells was highly variable among the specimens (Fig. S2C) per our annotation based on graphical clustering patterns and canonical cell-type specific markers for tumor epithelial/urothelial cells (*EPCAM*, *KRT7*), immune cells (*PTPRC*), stromal cells (*DCN*, *ACTA2*), and endothelial cells (*SELE*) (Fig. S3A).

We focused our analysis on tumor cell biology by subsetting and re-clustering the tumor epithelial cells from the main dataset (Fig. 1A). We excluded three tumors that did not meet a threshold of 150 tumor epithelial cells for analysis (UC04, VAR10, VAR11). Although neuroendocrine tumors are generally considered non-urothelial cancers,^2^ we included the tumor with small cell HV (VAR09) due to the presence of urothelial components within the tumor (carcinoma in situ and micropapillary variant). The final tumor epithelial dataset thus included three pure UCs (UC01-UC03) and nine HVs (VAR01-VAR09). To confirm the tumor content in this dataset, we used InferCNV to estimate the copy number profiles of all epithelial cell clusters using stromal and immune cells as reference (Fig. S3B).

**Figure 1.**
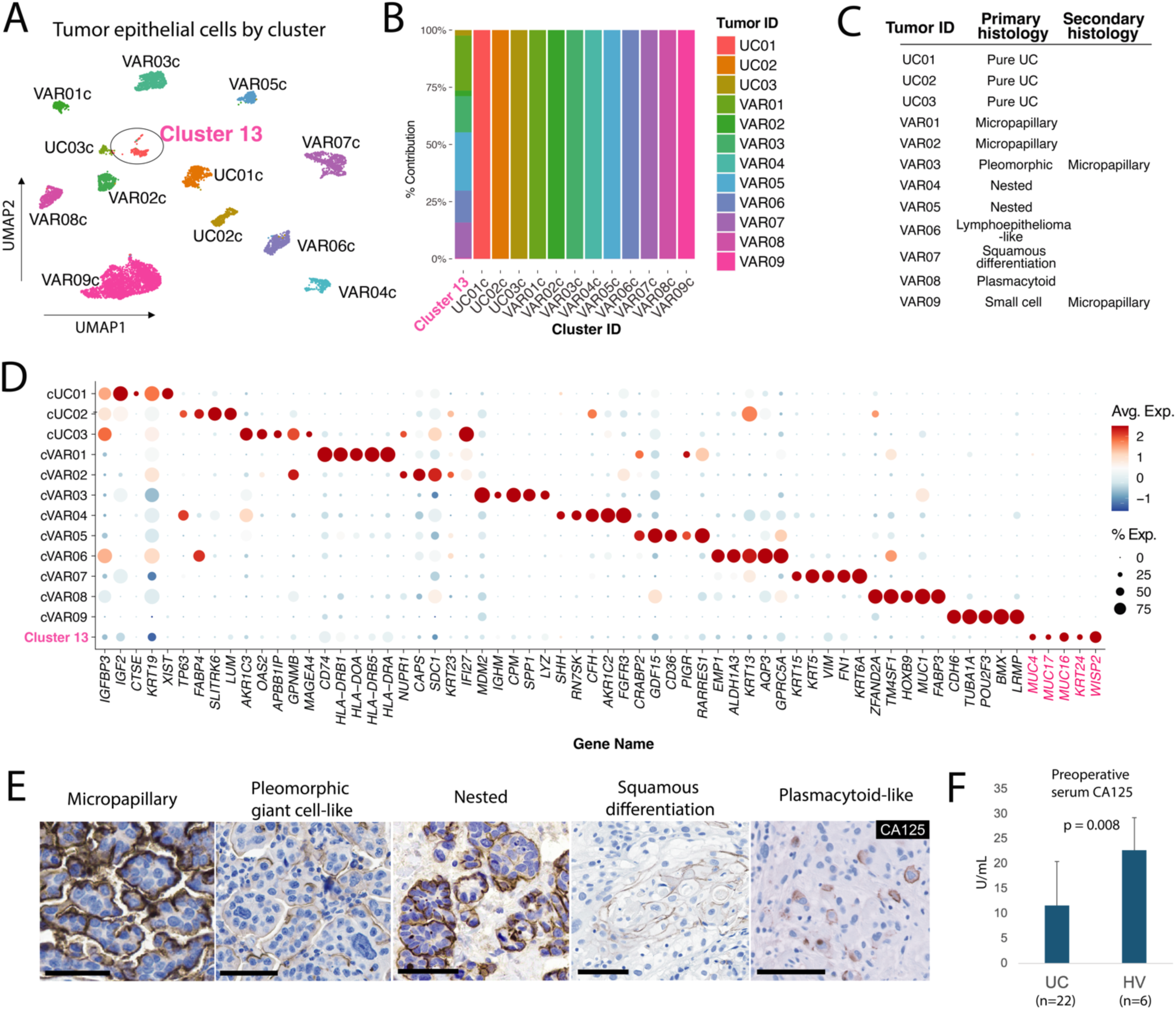
Top level clustering analysis of tumor epithelial cells and characterization of a common tumor cluster. (A) Clustering UMAPs of tumor epithelial cells (N = 8,553) extracted from the main dataset color-coded by cluster and annotated according to tumor ID. Cluster 13 (ellipse) is annotated separately due to contributions from multiple tumors. (B) Barchart of cluster composition by patient/tumor. (C) Table displaying primary and secondary histologic patterns observed in each tumor. (D) Curated dot plot of top differentially expressed genes (DEGs) by tumor cluster. (E) Immunohistochemistry of CA125 in primary tumor cells (VAR05) and nodal metastases (VAR09). Scale bar = 50 µm. (F) Preoperative serum CA125 values in patients with UC and HV bladder tumors.

Most tumor cells formed their own clusters corresponding to the tumor of origin and were named accordingly, i.e. VAR01c is the predominant cluster obtained from the VAR01 tumor (Fig. 1A). Interestingly, one cluster, which we named “Cluster 13” based on the number assigned by the clustering algorithm, was comprised of cells from multiple HV tumors (Fig. 1B-C). Differentially expressed genes (DEGs) for each tumor cluster were computed and curated, and *MUC16, WISP2, KRT24, MUC17,* and *MUC4* were among the top DEGs for Cluster 13 (Fig. 1D). To validate the existence of Cluster 13 cells histologically, we performed immunostaining of CA125 (encoded by *MUC16*) in HV (N=14) and UC (high-grade invasive and carcinoma in situ) tumors (N=20). We found a subpopulation of CA125+ cells in a variety of HV tumors with different subtypes (13/14) (Fig. 1E) but rarely in tumors with UC (1/11) or carcinoma in situ (CIS) histology (1/9). In tumors with mixed HV and UC components such as VAR03 and VAR05, CA125+ cells were present in the HV regions (Fig. 1E, pleomorphic giant cell-like, nested) but absent in the high-grade UC and CIS regions (Fig. S4). We did not detect the Cluster 13 signature or expression of *MUC16*, *KRT24*, and *WISP2* in a previously published bladder cancer scRNA-seq dataset derived from UC bladder tumors (Chen et al, Fig. S5A-B).^22^ Our results suggest that the cancer cells found in Cluster 13 represent a tumor cell state highly specific to, but not exclusive to, HV-containing tumors. To explore whether CA125 expression in these cells could be useful as a clinical biomarker, we prospectively assayed preoperative serum CA125 levels in bladder cancer patients undergoing surgery and found CA125 levels to be higher in those with HV components in their final pathology compared to those with UC only (22.7 ± 6.6 U/mL vs 11.6 ± 8.8 U/mL, p = 0.008) (Fig. 1F).

### Cluster 13 cells exhibit hallmarks of transcriptional convergence

To investigate the overall transcriptomic relationship among tumor clusters and to test whether similar HV subtypes share gene expression programs (e.g. micropapillary to micropapillary, nested to nested), an unsupervised partition-based graphical abstraction (PAGA) graph was generated. While we did not discern any prominent subtype-specific associations, we found that Cluster 13 cells formed a central node with an association to almost every other tumor cluster, even to those whose parent tumor did not contribute any cells to Cluster 13 (Fig. 2A). This result raised the possibility that Cluster 13 represents either a convergent cell state or a common progenitor cell state.

**Figure 2.**
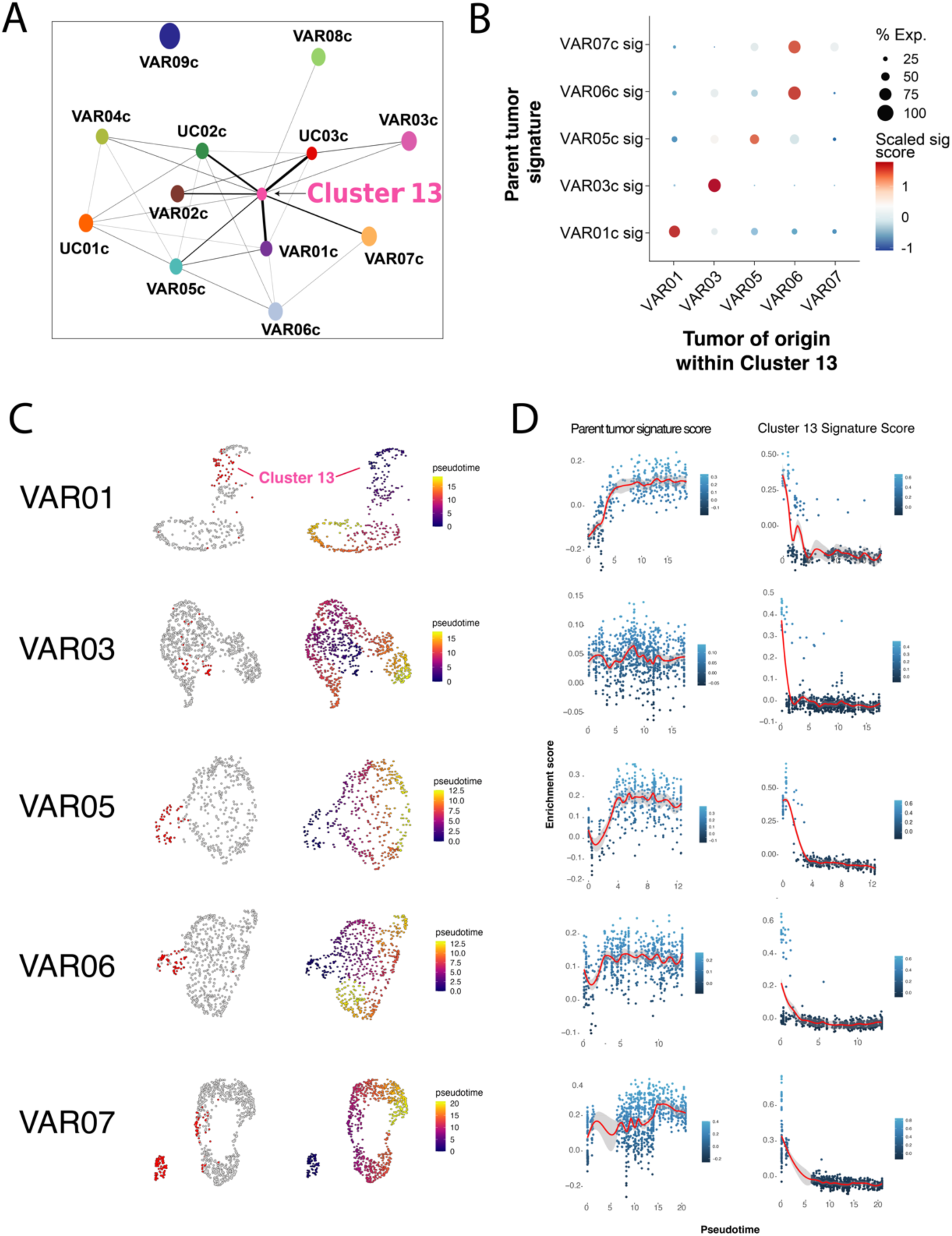
Transcriptional relationship between Cluster 13 and parent tumor cells. (A) Partition-based graphical abstraction of tumor cell clusters. (B) Dot plot of tumor signature scores relative to Cluster13 tumors of origin. (C) UMAP of individual tumors color-coded by Cluster 13 cells (red) and pseudotime using Cluster 13 cells as the starting point. (D) Expression along the pseudotime of Cluster 13 and parent tumor DEGs.

We thus sought to infer the relationship between the Cluster 13 cells and the parent tumor cells. We found that Cluster 13 cells bear the signature of the parent tumor with a high degree of specificity, supporting the likelihood that all cells within these tumors are clonally related (Fig. 2B). We performed pseudotime analysis for VAR01, VAR03, VAR05, VAR06, and VAR07 (Fig. 2C). The Cluster 13 cells were arbitrarily selected as the starting point for the pseudotime trajectory in each tumor to evaluate the relative signature enrichment between Cluster 13 cells and parent tumor cells (Fig. 3C). The Cluster 13 signature was anticorrelated with the parent tumor signature in four of five tumors (Fig. 3D), and the marked contrast of the Cluster 13 signature along the pseudotime in all five tumors suggests that Cluster 13 arises as a derivative of the parent tumor rather than vice versa. To further test the possibility that Cluster 13 is a progenitor cell state rather than a derivative tumor cell state, we generated a nine-gene bladder stem cell signature (*PROM1* (CD133), *POU5F1* (Oct4), *SOX2*, *ALDH1A1*, *SOX4*, *EZH2*, *YAP1*, *CD44*, and *KRT14*) based on previous studies in bladder cancer stem cells;^23–25^ and found no significant enrichment of this signature in Cluster 13 cells (Fig. S6). While scRNAseq alone cannot prove the temporal relationship between these cells, our results support the idea that cancer cells found in Cluster 13 are a convergent cell state in HV tumors.

**Figure 3.**
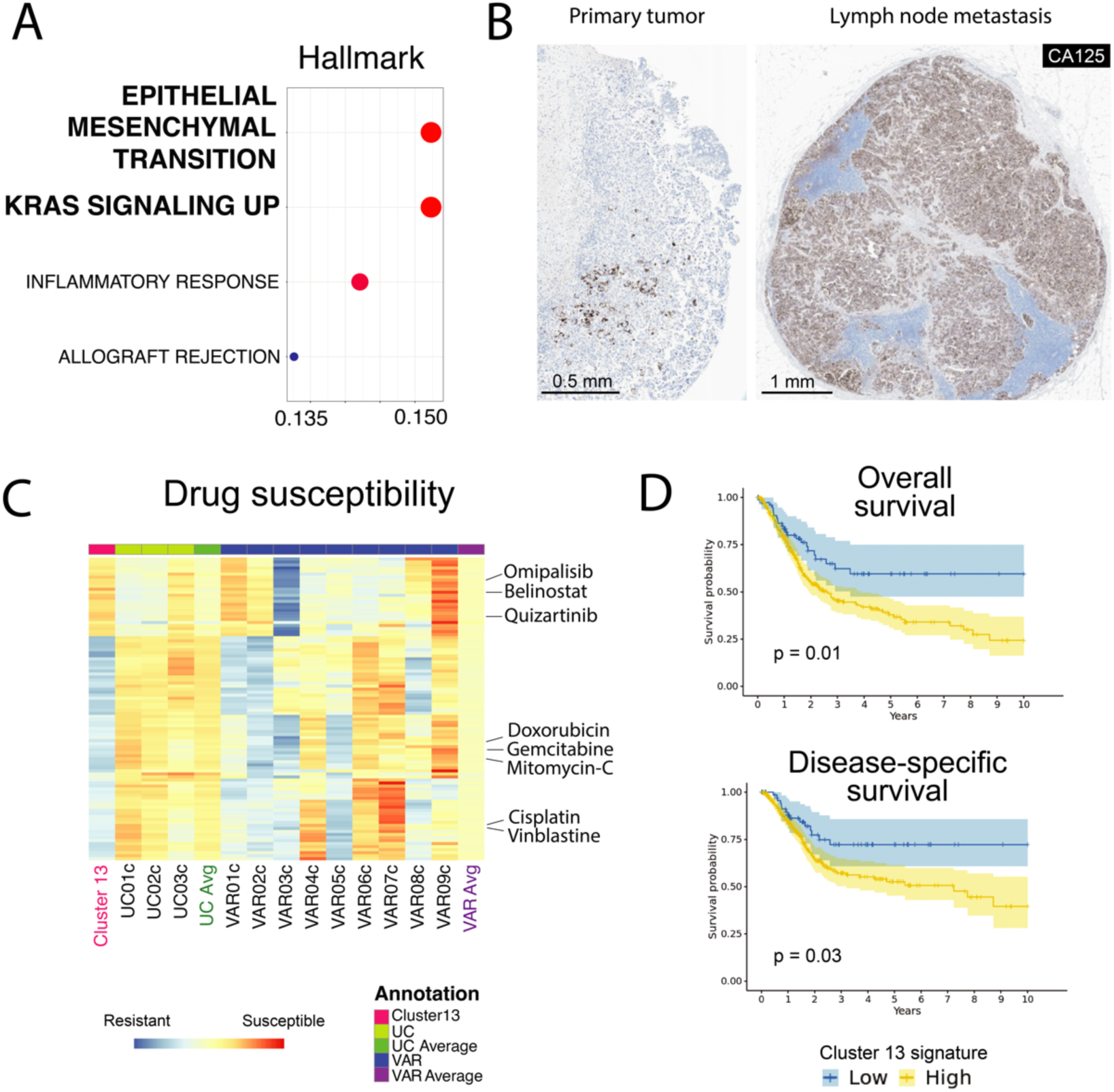
Cluster 13 is associated with metastasis and chemotherapeutic resistance. (A) Gene ontology analysis of Cluster 13 gene signature. (B) CA125 immunohistochemistry in a primary HV bladder tumor and the corresponding lymph node metastasis. (C) Drug susceptibility heatmap for gene signature individual tumor clusters and average UC and HV profiles. (D) Kaplan-Meier curves of overall and disease-specific survival according to Cluster 13 signature enrichment in TCGA-BLCA.

### Cluster 13 cells harbor adverse molecular features

Gene ontology (GO) analysis was performed on the DEGs for Cluster 13 cells and revealed a significant enrichment in epithelial-to-mesenchymal transition (EMT) and *KRAS* signaling gene sets (Fig. 3A). These findings raised the possibility that the Cluster 13 cells have more aggressive metastatic potential compared to non-Cluster 13 cells. Using CA125 again as a putative marker for Cluster 13 cells, we examined CA125 staining in five HV tumors with lymph node metastases and observed a striking homogeneous enrichment of CA125+ cells in the lymph nodes compared to the primary tumor in 4 of 5 cases (Fig. 3B).

We next evaluated the susceptibility of Cluster 13 cells to chemotherapy and targeted agents *in silico*. By training a drug response model using the Cancer Drug Response prediction using a Recommender System (CaDRReS) based on the Cancer Cell Line Encyclopedia (CCLE) database and Genomics of Drug Sensitivity in Cancer (GDSC) database, the estimated efficiency (percentage of tumor cells killed) for drugs from the GDSC database was inferred for each tumor cluster in our scRNA-seq dataset.^26^ Our analyses revealed that Cluster 13 cells were predicted to be more resistant to most chemotherapeutic agents, particularly in the case of conventional bladder cancer agents such as cisplatin, gemcitabine, and mitomycin C, compared to UC and non-Cluster 13 HV cells (Fig. 3C). Consistent with these adverse features, tumors that harbor higher Cluster 13 signature scores in TCGA-BLCA had worse overall survival and disease-specific survival (Fig. 3D).

Taken together, these results indicate that HV tumors contain a cancer cell state that is enriched in metastases and are predicted to be more resistant to chemotherapy. This cell state offers a potential mechanism to help explain why HV tumors are more aggressive than UC tumors.

### Histologic variants exhibit transcriptional hallmarks of histologically similar non-urothelial cell types

Our analyses enabled us to examine whether HVs share molecular features with other histologically similar but non-urothelial cell types. Specifically, we investigated this possibility in tumor cells from VAR09 (small cell) and VAR08 (plasmacytoid), which exhibited low enrichment of urothelial differentiation genes (Fig. S7A-B).

It has been proposed that small cell bladder cancers (SCBC) exhibit similarities with small cell lung cancers (SCLC) based on similar genomic alterations, but transcriptomic evidence has been limited.^27^ For VAR09, we applied a molecular subtyping schema based on the expression of *ASCL1*, *NEUROD1*, *POU2F3*, and *YAP1* to the VAR09 tumor cells and noted enrichment in *POU2F3*, a gene associated with the SCLC-P subtype, or the “tuft cell-like variant” (Fig. 4A).^28–30^ *POU2F3* expression was highly expressed throughout the VAR09 tumor cells along with downstream targets *AVIL*, *SOX9*, and *PTGS1*, expression of which was specific to VAR09 compared to other HVs. (Fig. 4B).

**Figure 4.**
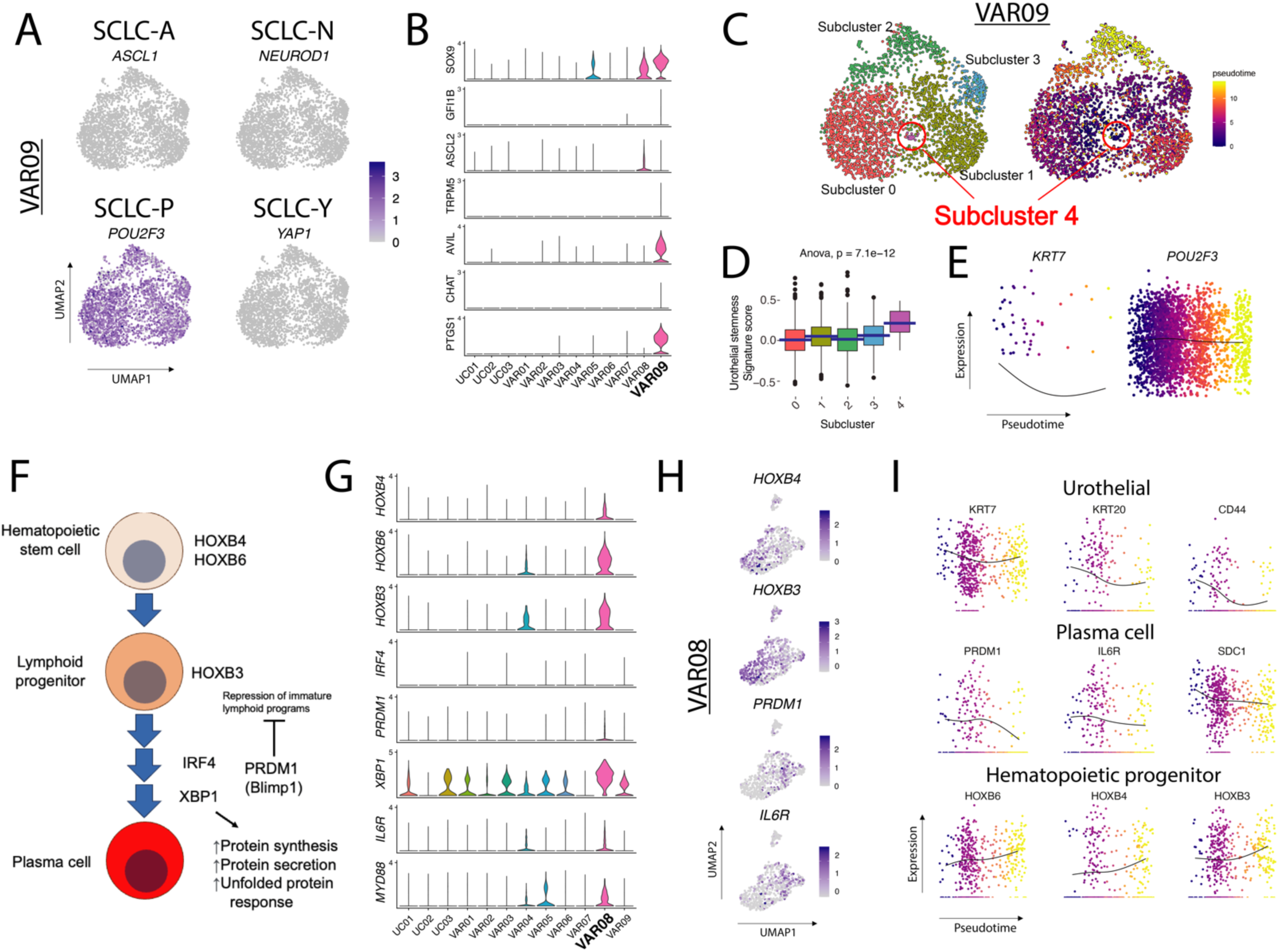
Nonurothelial transcriptional programs in VAR09 and VAR08. (A) Feature plots of small cell lung cancer (SCLC) molecular subtype-defining genes ASCL1 (SCLC-A), NEUROD1 (SCLC-N), POU2F3 (SCLC-P), and YAP1 (SCLC-Y) expression in VAR09. (B) Expression of POU2F3 downstream targets across tumor clusters. (C) UMAP of VAR09 color-coded by subcluster and pseudotime using subcluster 4 as starting point. (D) Urothelial stemness signature score among VAR09 subclusters. (E) Expression of KRT7 and POU2F3 along the VAR09 pseudotime. (F) Schematic of HOXB genes and transcription factors IRF4, PRDM1, and XBP1 along the plasma cell lineage. (G) Expression of plasma cell-related genes among tumor clusters. (H) Feature plots of HOXB4, HOXB3, PRDM1, and IL6R expression in VAR08 cells. (I) Expression of urothelial and lymphoid genes along the VAR08 pseudotime.

Although most VAR09 cells lacked expression of canonical urothelial markers, *KRT7* expression was detected and was primarily localized to subcluster 4 (Fig. 4C). This cluster also harbored the highest stemness signature (Fig. 4D), so we performed pseudotime analysis using cells from subcluster 4 as a starting point, which showed a decrease in *KRT7* expression along the pseudotime trajectory while *POU2F3* largely remained constant (Fig. 4C, 4E). The coexistence of a *KRT7+* and *POU2F3+* cluster with progenitor-like features supports the hypothesis that small cell bladder cancer cells arise from a urothelial origin despite the low expression of urothelial genes.^27^

Due to their plasmacytoid appearance, we next investigated whether tumor cells from VAR08 exhibited hallmarks of hematopoiesis and plasma cell differentiation. We noted that HOX genes, transcription factors important for hematopoiesis that are known to be upregulated in some bladder cancers, were among the top DEGs for VAR08 (Fig. 2D).^31–35^ HOXB genes, which are required for hematopoietic stem cell (HSC) maintenance (*HOXB4*, *HOXB6*) and B-cell maturation (*HOXB3*), were enriched in VAR08 compared to other tumors (Fig. S8).^36–39^ Immune cell signatures derived from our scRNA-seq dataset were generated (Myeloid, T-cell, B-cell, and Plasma cell) (Fig. S9A-B), and we found that VAR08 cells were enriched for the plasma cell signature (Fig. S9C). Expression of plasma cell-specific transcription factors *PRDM1* and *XBP1* and surface marker IL6R was elevated in VAR08 tumor cells, although the upstream activator *IRF4* was notably absent (Fig. S10A).^40^ *MYD88*, a gene associated with lymphoplasmacytic lymphomas, was also detected in VAR08 tumor cells.^41^ The major determinants of plasma cell differentiation harbored within VAR08 were thus identified (Fig. 4F-G).^42^ Protein chaperones (*HSPA1B*, *HSPA5*) and protein synthesis genes (*ELL2*, *EIF2AK3*) were also highly expressed in tumor cells from VAR08 (Fig. S10A), and gene sets related to the unfolded protein response and protein secretion were also enriched (Fig. S10B). These features are consistent with the upregulation of downstream targets of *PRDM1* and *XBP1* similar to the transcriptional programs found in plasma cells.^43,44^

Next, we asked whether different stages of plasma cell maturation could be observed in this tumor. We observed that *HOXB4* and *HOXB3*, typically expressed earlier in the lineage, were anticorrelated with late genes IL6R and *PRDM1*, a broad repressor of immature transcriptional programs (Fig. 4H).^44,45^ This suggested the coexistence of a HSC-like state (HOX^high^) along with a more differentiated plasma-cell like state (*PRDM1*^high^/*IL6R*^high^). When we performed pseudotime analysis starting from VAR08 cells with highest *KRT7* expression, a surrogate for urothelial differentiation, we observed a rise in HOX gene expression and a concomitant fall in the expression of *KRT20*, *CD44*, and *SDC1* (CD138) (Fig. 4I) along the pseudotime trajectory, indicating that VAR08 tumor cells may transition from a plasma cell-like urothelial state towards a more dedifferentiated HSC-like state. Of note, neither plasma cell lineage (*CD34*, *PTPRC* (CD45), *CD19*, *MS4A1* (CD20), *CD27, CD38*) nor immunoglobulin gene expression was detected in VAR08, suggesting that activation of hematopoietic transcriptional programs in urothelial cells does not necessarily result in expression of hematopoietic surface lineage markers.

### TM4SF1 is a surface protein broadly enriched in histologic variant tumor cells

Having identified and characterized the Cluster 13 cell state in HVs, we next asked whether our scRNA-seq results could help identify any molecular features broadly enriched in HV tumor cells compared to UCs; defining such features would facilitate the development of HV-specific targeted therapies. We categorized all tumor cells as HV or UC according to the histology of the parent tumor and computed the DEGs (Fig. 5A). *TM4SF1*, a gene implicated in bladder cancer as a cell cycle and apoptosis regulator, was the top DEG in the HV group.^26–28^ Most HV tumor clusters, including Cluster 13, exhibited higher expression of *TM4SF1* compared to those from pure UC tumors (Fig. 5B). VAR02c and VAR09c were the only tumor cell clusters with absent *TM4SF1* expression.

**Figure 5.**
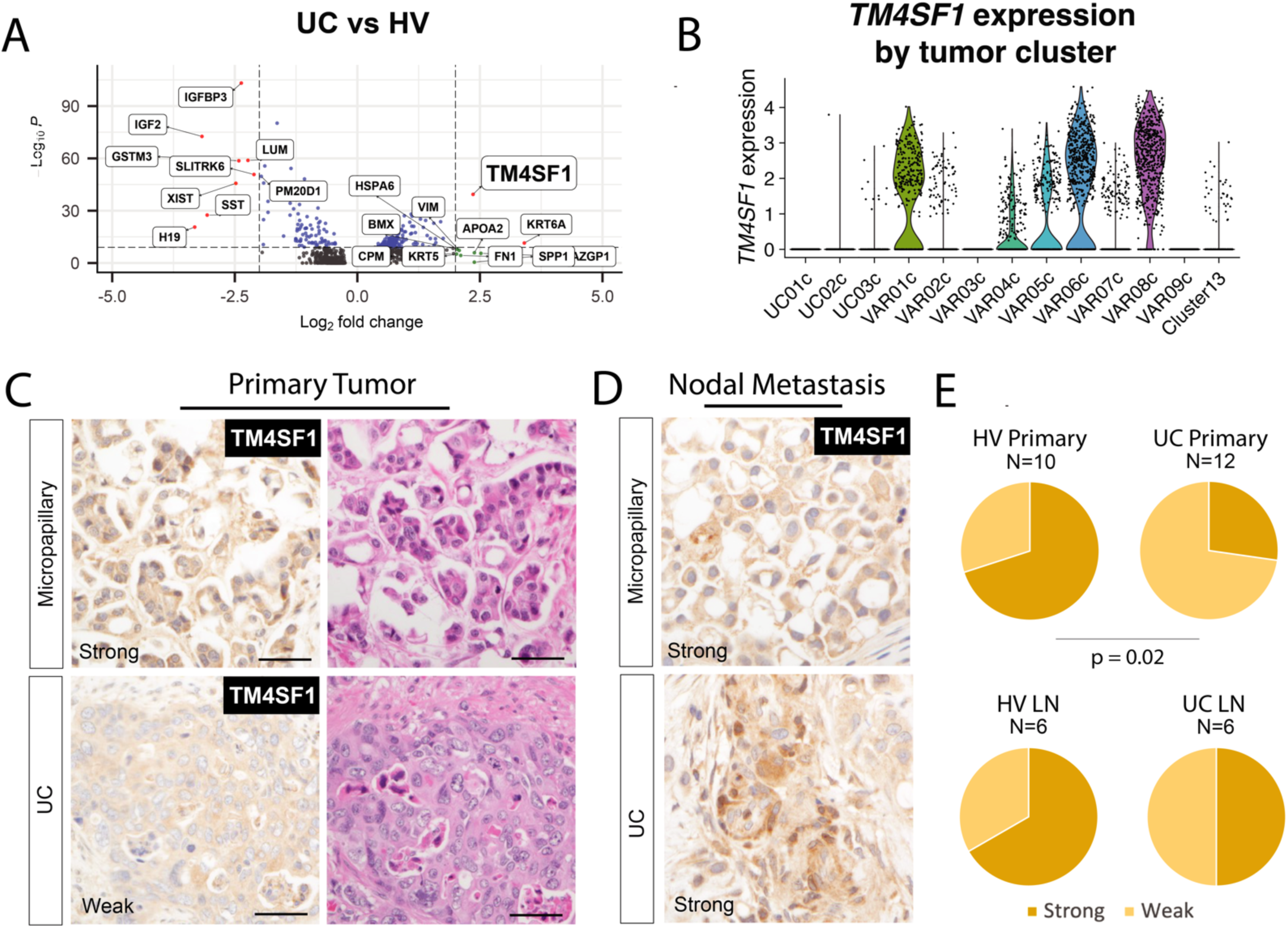
Identification of TM4SF1 as a gene enriched in HVs. (A) Volcano plot comparison of all UC and HV cells after downsampling (N = 150 per patient). (B) Violin plots of TM4SF1 expression by tumor cluster. (C) Correlation plots of TM4SF1 and EMP1, EZR, CLDN4, and KRT19. (D-E) Immunohistochemistry of TM4SF1 in a validation cohort of HV and UC (D) primary tumors and (E) lymph node metastases. (F) Semiquantitative comparison of TM4SF1 staining in HV and UCs.

Consistent with previous reports, we confirmed that high *TM4SF1* expression is associated with basal tumor signatures (Fig. S11A) and adverse clinical outcomes in TCGA-BLCA (Fig. S11B-C).^46^ In our tumor epithelial data set, genes with the strongest positive correlation with *TM4SF1* expression within the HV tumor cells were *EMP1*, *CLDN4*, *EZR*, and *KRT19* (Fig. S12A-B). We checked the associations within each *TM4SF1*-expressing tumor in our scRNA-seq dataset and found these to be positive and statistically significant in each case (Fig. S12C). *EMP1*, a gene implicated in cisplatin resistance and cancer recurrence,^47–49^ and *CLDN4*, a tight junction gene implicated in facilitating aggressive biology in bladder cancer,^50^ were also positively associated with *TM4SF1* in TCGA-BLCA (Fig. S12D). Interestingly, we did not observe a statistically significant association between *TM4SF1* expression and *SOX2*, *DDR1*, *MMP2*, or *MMP9* expression (data not shown), suggesting that the expression of *TM4SF1* in HVs may be regulated differently than what has been previously described in cell lines and nonurothelial cancers.^51,52^

Using immunohistochemistry, we validated TM4SF1 protein expression in HV and UC cells, both in primary tumors and lymph node metastases (Fig. 5C-D). Consistent with our sequencing results, quantification of TM4SF1 staining using a binary “strong” and “weak” scoring system (see methods) demonstrated more frequent strong staining in HV primary tumors compared to UC primary tumors (p = 0.02) (Fig. 5E).

### TM4SF1-CAR T cells demonstrate *in vitro* and *in vivo* activity against bladder cancer cells

The enrichment of *TM4SF1* expression in HVs and its cell surface expression made it a compelling candidate for developing a targeted therapeutic strategy. Expression of TM4SF1 is high across a number of tumor types, and its inverse correlation with *PVRL4* (NECTIN4) expression in TCGA-BLCA and CCLE (Fig. S12D, Fig. S13) suggests that TM4SF1-directed therapies might be complementary to enfortumab vedotin (EV) therapy, an antibody-drug conjugate that targets NECTIN4 that was recently approved for frontline treatment of patients with locally-advanced/metastatic urothelial cancers.^53,54^

Given that there are no FDA-approved TM4SF1-directed therapeutic agents, we next asked whether TM4SF1 could be targeted by chimeric antigen receptor (CAR) T cell therapy. To test this, we utilized a previously published TM4SF1 single-chain variable fragment (scFv) binder and incorporated this into a 41BB-based CAR bone in two configurations (VH-VL (CAR1) and VL-VH (CAR2)) (Fig. 6A). We tested both CAR T candidates against six bladder cancer cell lines with variable levels of endogenous *TM4SF1* mRNA expression and surface protein expression (Fig 6B). Whereas the TM4SF1-CAR T cells exhibited anti-tumor activity against bladder cancer lines expressing TM4SF1 (including UMUC3, T24, 5637, 253JBV and UMUC1), the TM4SF1-CAR T cells did not kill HT1376, which are negative for TM4SF1 (Fig. 6C). We also found CAR1 had slightly better activity *in vitro*. To validate the specificity of our CARs, we used CRISPR/Cas9 to generate *TM4SF1* knockouts (KO) in the UMUC3 cell line, which abolished the anti-tumor activity of TM4SF1-CAR T cells (Fig. S14A-B).

**Figure 6.**
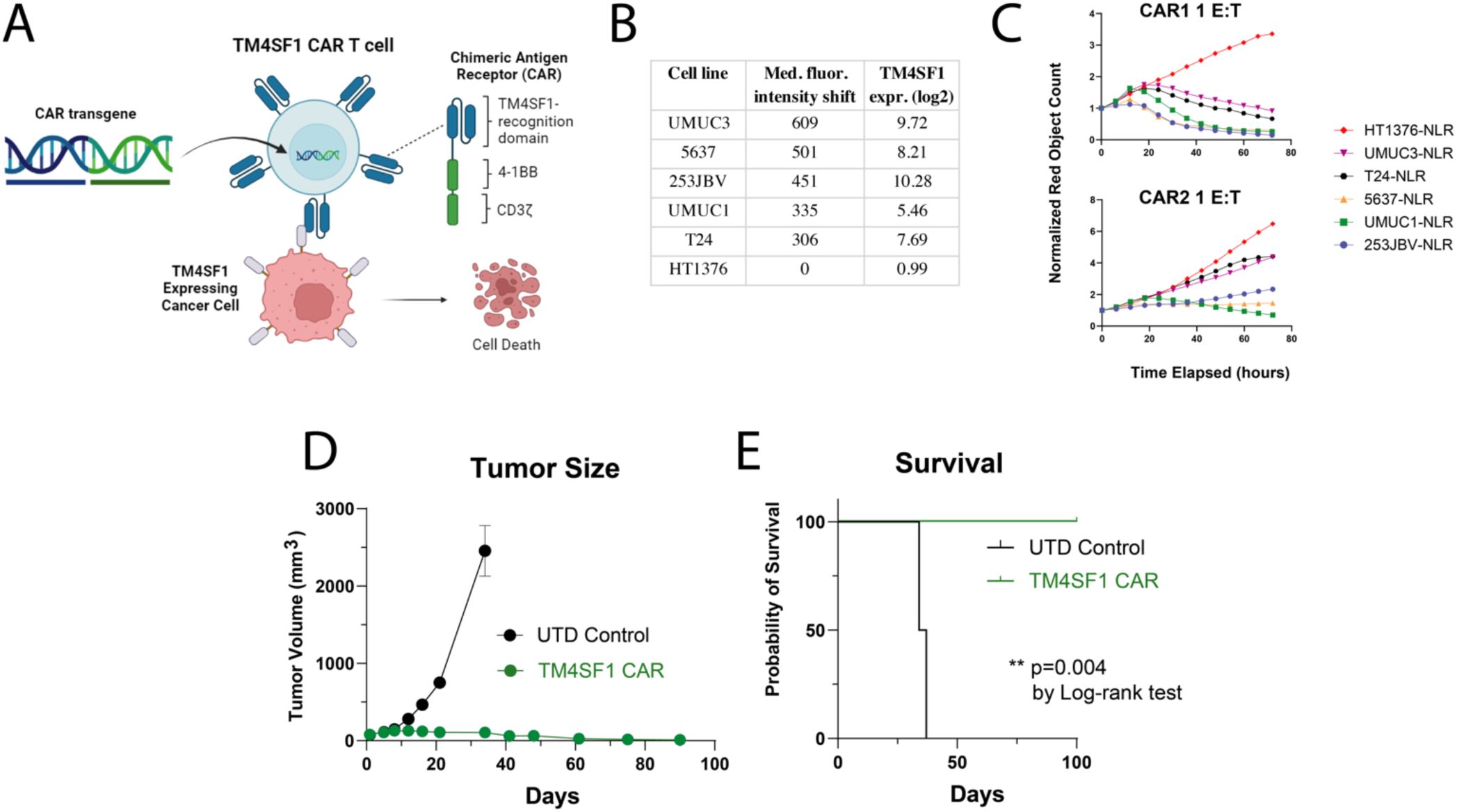
Efficacy of TM4SF1 CAR T cells *in vivo* and *in vitro*. (A) Schematic for generating TM4SF1-CAR T cells. (B) Bladder cancer cell lines and TM4SF1 expression determined by flow cytometric fluorescent antibody detection and mRNA expression. (C) Quantification of *in vitro* TM4SF1-CAR1 and CAR2 activity against bladder cancer cell lines using IncuCyte co-culture assay with 1:1 effector:target cell ratio. (D) Schematic for xenograft generation from the UMUC3 cell line and *in vivo* TM4SF1-CAR1 testing. (E) Tumor size comparisons between TM4SF1-CAR treated and untreated mice. (F) Kaplan-Meier survival analysis of treated and untreated mice.

Finally, we tested CAR1 against xenografts derived from the UMUC3 cell line (Fig. S15), which was selected for its high TM4SF1 expression and absent NECTIN4 expression. We found that CAR1 exhibited potent anti-tumor activity against these tumors *in vivo*. Whereas control mice all died by day 37, mice treated with TM4SF1-CAR1 cells had complete and durable responses, even up to day 100 (n=5 mice, Fig. 6D-E). Importantly, mice treated with TM4SF1-CAR1 cells had stable weights (Fig. S16) and no overt pulmonary toxicity. Taken together, these data demonstrate that TM4SF1 could be a new therapeutic target for HV bladder cancers, including tumors lacking *NECTIN4* expression, and can be successfully targeted using CAR T cell therapy.

## DISCUSSION

In this study, we demonstrate that scRNA-seq can be used to identify molecular features for rare, understudied cancer types such as HV bladder cancers. Here, we have described a novel cancer cell state (Cluster 13) with clinical and mechanistic significance and a targetable protein (*TM4SF1*) in HV bladder cancers. As HVs are poorly understood in part because they are heterogenous and uncommon, scRNAseq enabled us to derive insights about HV cancer biology in a relatively small cohort of tumors. Our study underscores the potential of scRNAseq technologies in precision cancer medicine.^55^

The identification of a distinct “Cluster 13” cell state, which was found in more than half of the sequenced HV tumors and can be detected using *MUC16* (CA125) as a marker, has potentially important clinical implications for HV bladder cancers. Since CA125+ cells are found in most HVs and are enriched in metastatic disease, a deeper characterization of this cell state may lead to new unified strategies to treat tumors that otherwise exhibit a great degree of heterogeneity. Although tumor cells harboring this cell state are predicted to be more resistant to conventional chemotherapeutics used for bladder cancer such as cisplatin, gemcitabine, doxorubicin, vinblastine, and mitomycin C, several United States Food and Drug Administration (FDA)-approved agents including omipalisib (PI3K/mTOR inhibitor), belinostat (histone deacetylase inhibitor), and quizartinib (FLT3 inhibitor) were predicted to be more effective against this group of cells (Cluster 13) compared to other tumor cells.

The specific expression of *MUC16* (CA125) and other mucin genes in this cell state is intriguing. CA125, a well described gene more commonly associated with ovarian and pancreatic cancers,^56–59^ is a membrane-bound mucin protein that can promote cancer invasion and metastasis, and it has also been associated with therapeutic resistance in bladder cancer.^60–62^ It will be important to establish in future studies how CA125 contributes to HV biology and clinical behavior. Serum CA125 levels have long been used for the clinical surveillance of ovarian cancer and may have diagnostic and prognostic implications for other cancers.^63,64^ Here we show that patients with HV tumors have higher serum CA125 levels compared to patients with UC tumors, supporting its potential use as a biomarker in bladder cancer and could be useful for serological monitoring of HV tumors.

The origin of the cancer cell state identified in Cluster 13 remains an important question. While our data suggest that the Cluster 13 cell state is a shared state that is found in different HV tumors, the temporal relationship between Cluster 13 cells and other cancer cells within each tumor cannot be determined using scRNA-seq alone. It remains possible that the Cluster 13 cell state represents a common precursor cell state for HV tumors. The existence of a common cell state that has metastatic potential and is chemotherapy-resistant among diverse HV tumors suggests that a common mechanism may underlie their aggressive behavior. Investigating how this cell state behaves functionally and how it arises may help inform our understanding of bladder cancer tumor evolution and metastasis.

We also show that HV tumors exhibit transcriptional programs characteristic of the non-urothelial cell types to which they share histologic resemblance. This raises the possibility that HVs could be treated using agents targeting those other tumor cell types. Appropriating therapies designed for other cancers has been shown empirically to be effective in the case of SCBC and SCLC, and we now provide evidence for how SCBC and SCLC can have overlapping transcriptional programs. We additionally demonstrate the existence of plasma cell transcriptional programs in the plasmacytoid HV. This provides a rationale to test whether therapies designed for plasma cell neoplasms could be effective for this HV in future studies.

Our discovery that most HV tumors exhibit enriched expression of *TM4SF1*, a gene that encodes a surface protein that has already been implicated in the pathogenesis of aggressive bladder cancers and other cancer cell types, has therapeutic implications.^46,51,52,65^ *TM4SF1* is a promising target because its expression is not limited to HV bladder cancers and its negative association with *PVRL4/PRR4* (NECTIN4) expression suggests that targeted therapy against *TM4SF1* could complement existing targeted agents.

Antibody-mediated inhibition of *TM4SF1* has been previously shown to have therapeutic potential against cancer stem cells *in vitro*;^66^ we now demonstrate durable anti-tumor responses in mice bearing xenografts with minimal toxicity. Our preclinical testing of TM4SF1-CAR T cells thus lays a foundation for future clinical trials in bladder cancer and other tumor types expressing TM4SF1.

The primary limitation of our scRNA-seq dataset is the relatively low cell capture rate. While this is a known limitation of the Seq-well platform and there was variable quality and viability of tumor specimens collected during surgery, we had sufficient cell numbers to investigate tumor epithelial cells. Our ability to compare differences in the tumor microenvironment and identify intercellular interactions, however, was limited. To address the low sample size of sequenced tumors, we used an existing scRNA-seq dataset as well as the TCGA-BLCA dataset to supplement our analyses.

In conclusion, our study demonstrates that HV subtypes in bladder cancer harbor a clinically significant CA125+ cell state, express a surface antigen that is targetable using CAR T cells, and share transcriptional features with histologically similar non-urothelial cancers. These findings lay a foundation for further translational investigation into these rare, poorly understood tumors.

