## Supplemental Materials for "Histologic variants in bladder cancer harbor aggressive molecular features including TM4SF1 expression and a CA125+ cell state"

**Table S1. Clinical and pathologic characteristics of patients and sequenced bladder cancer tissues**

| ID | Age | Sex | Race | Neoadjuvant therapy | Operation | Stage | HV subtype 1 | HV subtype 2 | CIS present |
|---|---|---|---|---|---|---|---|---|---|
| UC01 | 60 | F | Hispanic | chemotherapy | cystectomy | T2 | | | no |
| UC02 | 74 | M | White | immunotherapy | cystectomy | Ta | | | no |
| UC03 | 71 | M | White | none | TURBT | T1 | | | no |
| UC04 | 73 | F | White | none | cystectomy | T1 | | | yes |
| VAR01 | 68 | M | White | none | cystectomy | T2 | micropapillary | | yes |
| VAR02 | 87 | M | White | chemotherapy | TURBT | T2 | micropapillary | | no |
| VAR03 | 57 | M | White | none | TURBT | T1 | pleomorphic giant cell-like | micropapillary | no |
| VAR04 | 87 | M | Asian | none | TURBT | T2 | nested | | no |
| VAR05 | 67 | M | White | chemotherapy | cystectomy | T4 | nested | | yes |
| VAR06 | 82 | M | White | none | cystectomy | T2 | lymphoepithelioma-like | | yes |
| VAR07 | 44 | M | Asian | chemotherapy | cystectomy | T2 | squamous differentiation | | yes |
| VAR08 | 60 | M | Pacific Islander | none | cystectomy | T4 | plasmacytoid | | no |
| VAR09 | 84 | F | White | chemotherapy | cystectomy | T3 | small cell | micropapillary | no |
| VAR10 | 67 | F | White | chemotherapy | cystectomy | T4 | pure squamous | | no |
| VAR11 | 61 | M | Other | chemotherapy | cystectomy | T4 | squamous differentiation | sarcomatoid differentiation | no |

**Figure S1. Representative H&E stains from each sequenced tumor.**

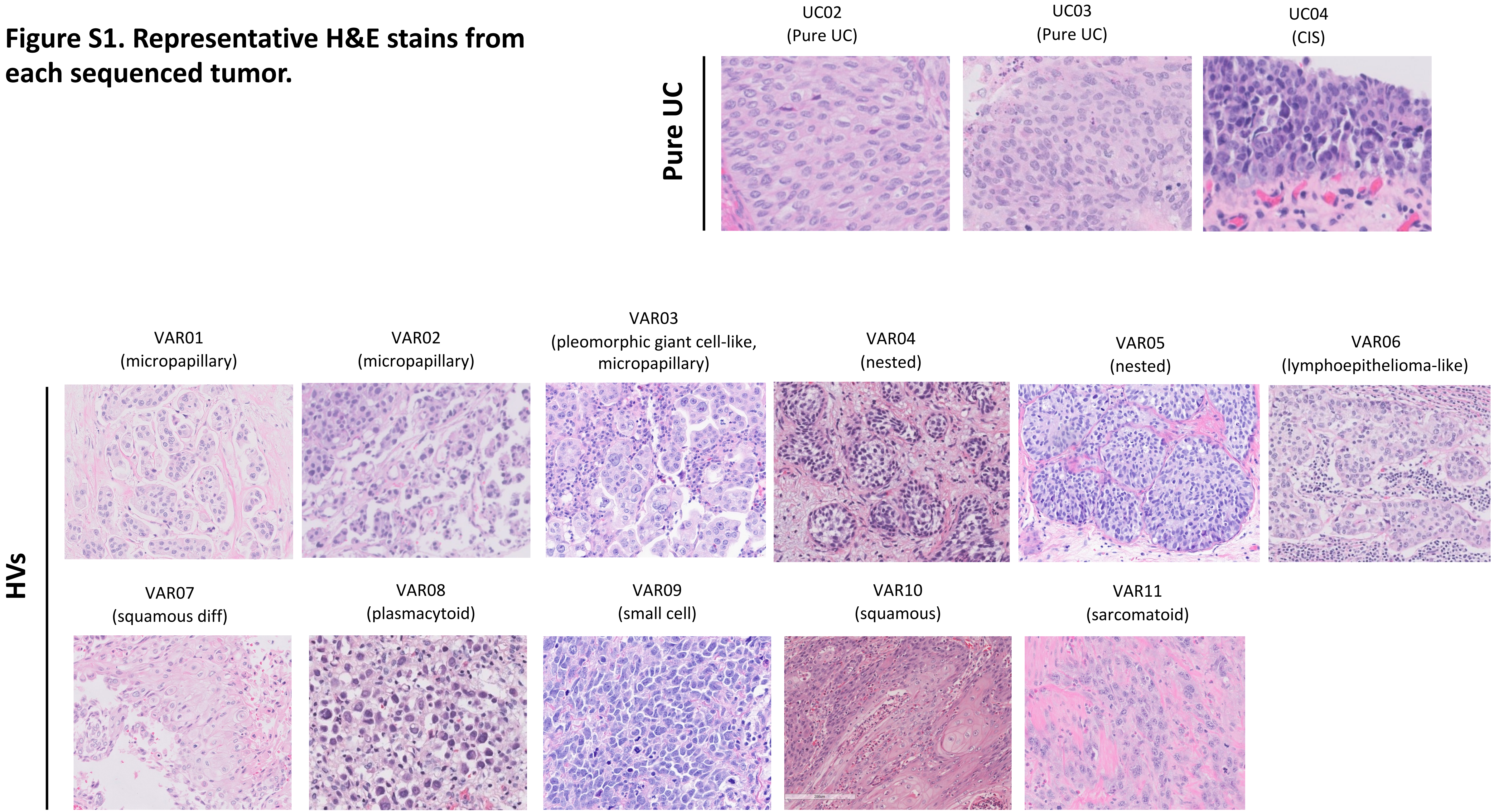

**Figure S2. Single cell dataset of variant and pure urothelial tumor epithelial cells.** (A) Tissue acquisition and scRNA-seq workflow for primary bladder tumors. (B) UMAP of full dataset color-coded by broad cell type. (C) Bar chart of cell counts obtained from each patient/tumor.

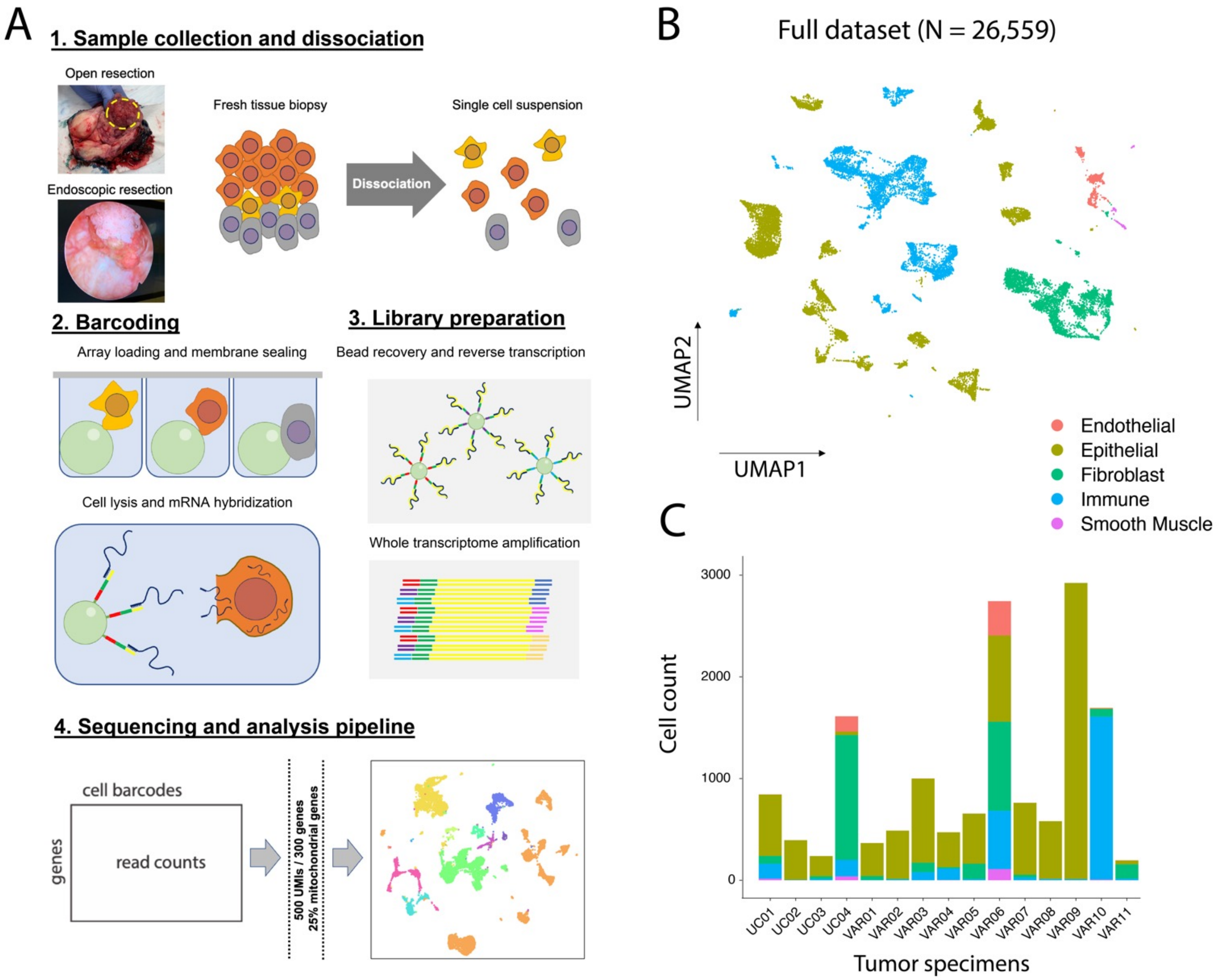

**Figure S3. Annotation of cell types and confirmation of tumor content.** (A) Feature plots of genes used to determine top-level annotations for epithelial cells (EPCAM, KRT7), immune cells (PTPRC), fibroblasts (DCN), smooth muscle (ACTA2), and endothelial cells (SELE). (B) InferCNV analysis of all tumor epithelial cells using tumor microenvironment components for comparison.

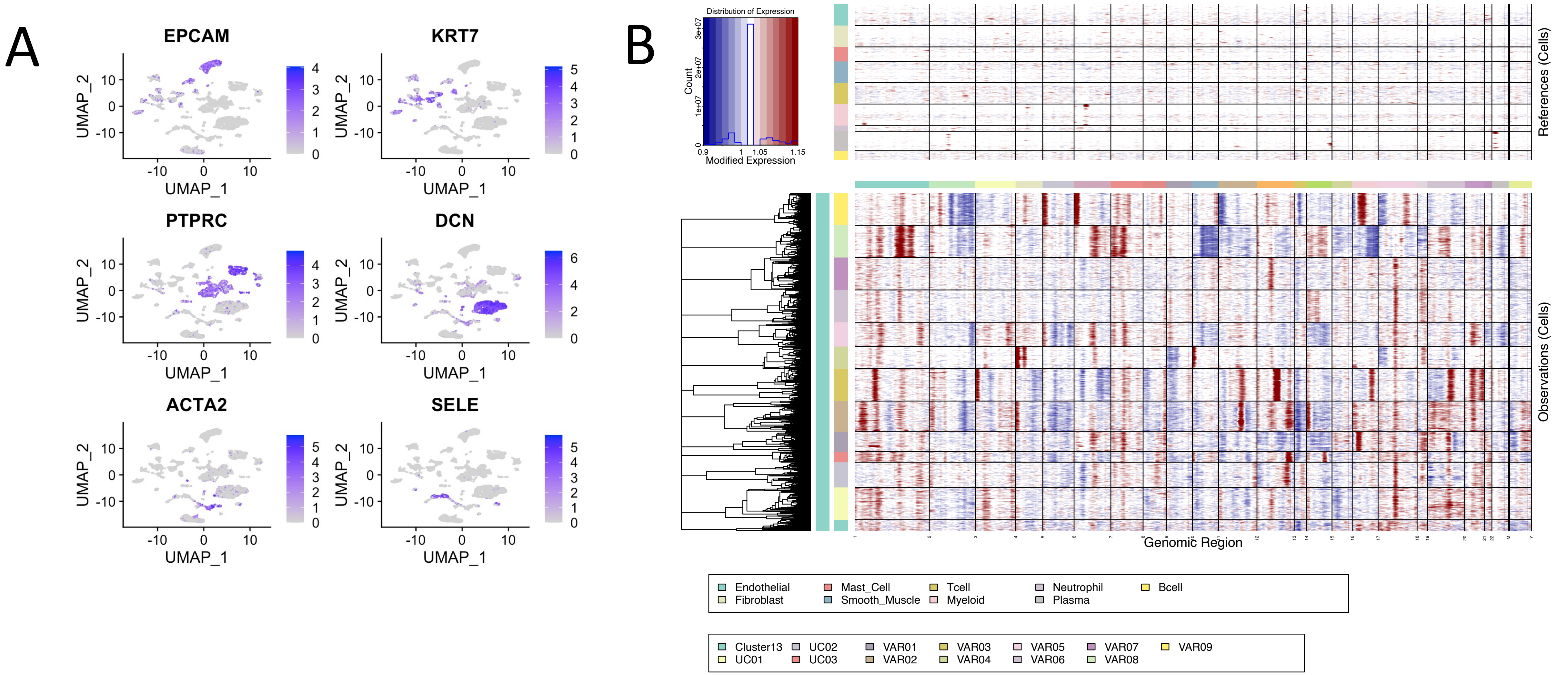

**Figure S4. Negative CA125 (MUC16) staining in variant and pure UC tumor components.** Scale bar = 50 μm.

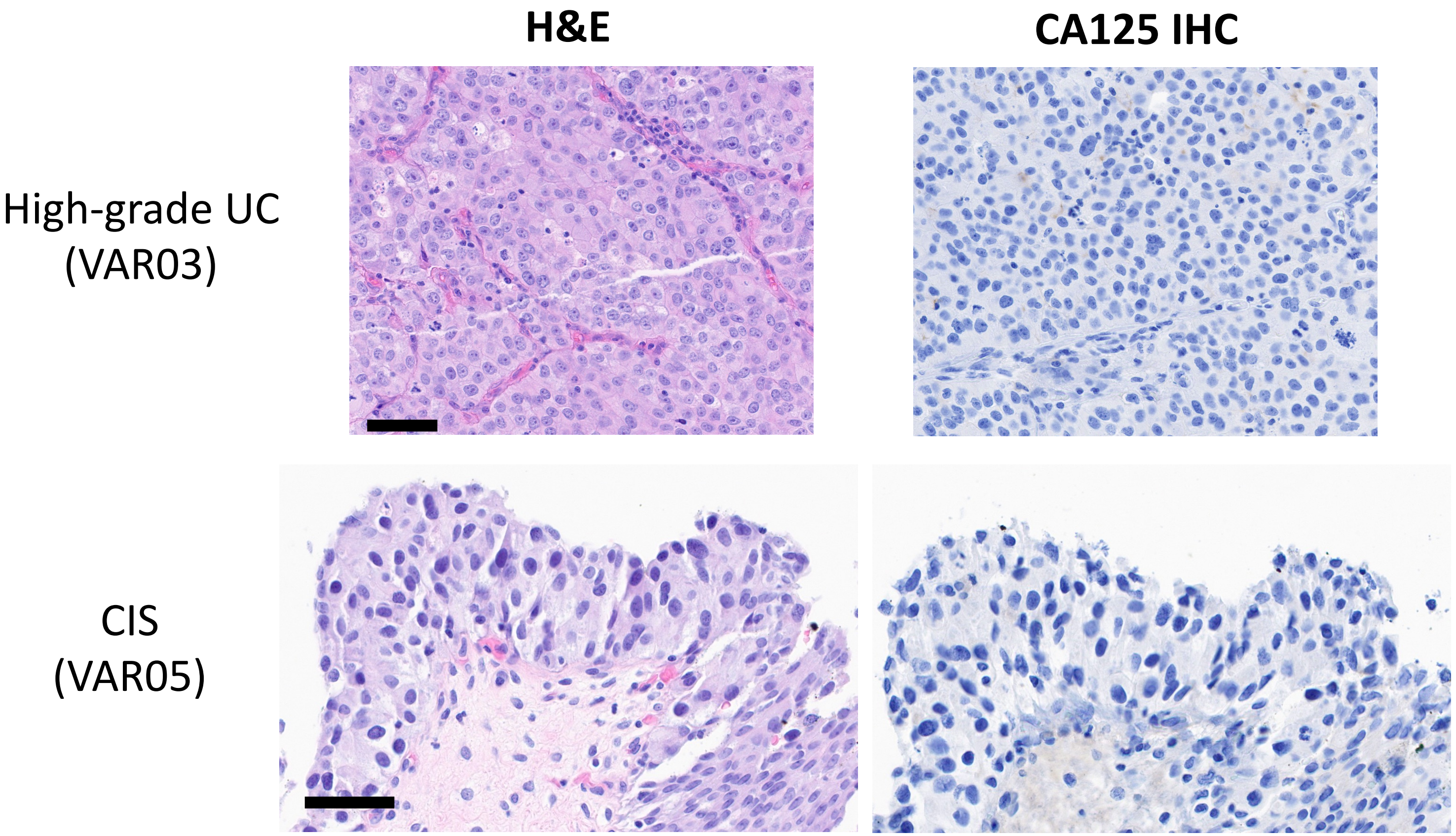

**Figure S5. Detection of Cluster 13 signature in an external scRNAseq data set (Chen et al).** (A) Feature plot of Cluster 13 signature gene set enrichment. (B) Feature plot demonstrating expression of individual Cluster 13-defining genes.

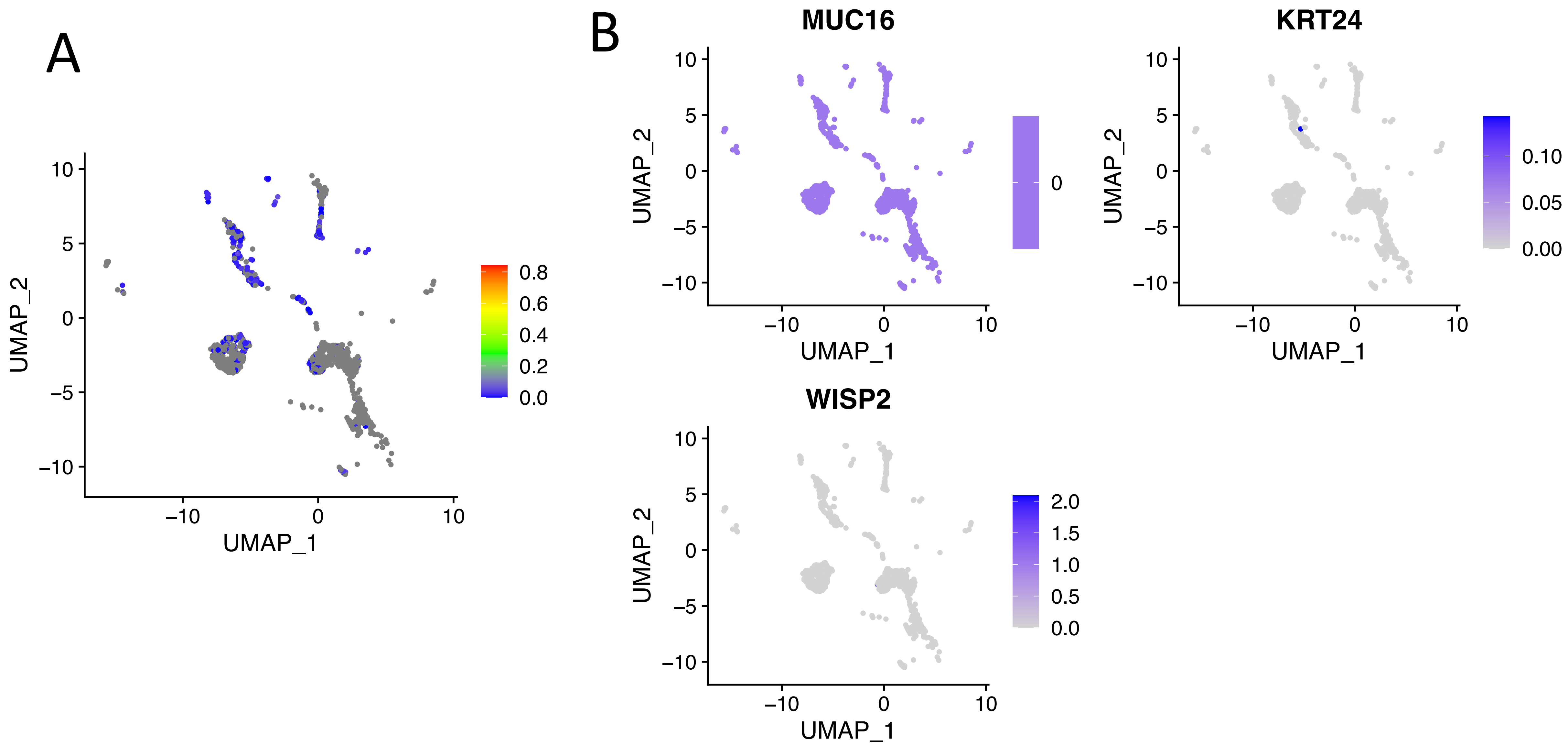

**Figure S6. Enrichment of bladder cancer stem cell signature (*PROM1* (CD133), *POU5F1* (Oct4), *SOX2*, *ALDH1A1*, *SOX4*, *EZH2*, *YAP1*, *CD44*, and *KRT14*) along the pseudotime for VAR01, VAR03, VAR05, VAR06, and VAR07.**

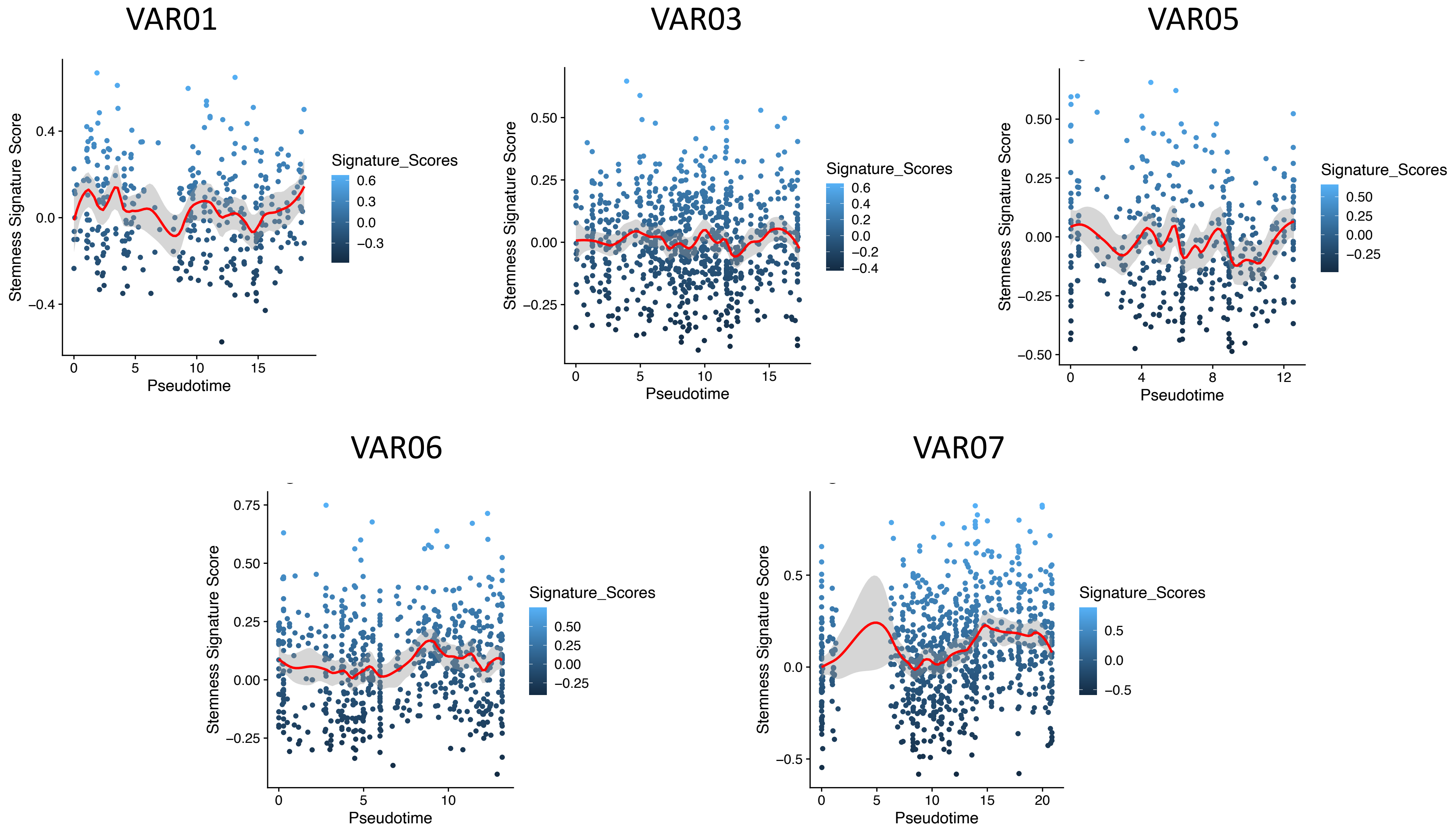

**Figure S7. Assessment of urothelial carcinoma gene signature in each tumor.** (A) Expression of urothelial carcinoma gene signature (Mo et al) in each tumor by individual gene. (B) Urothelial carcinoma signature enrichment in each tumor.

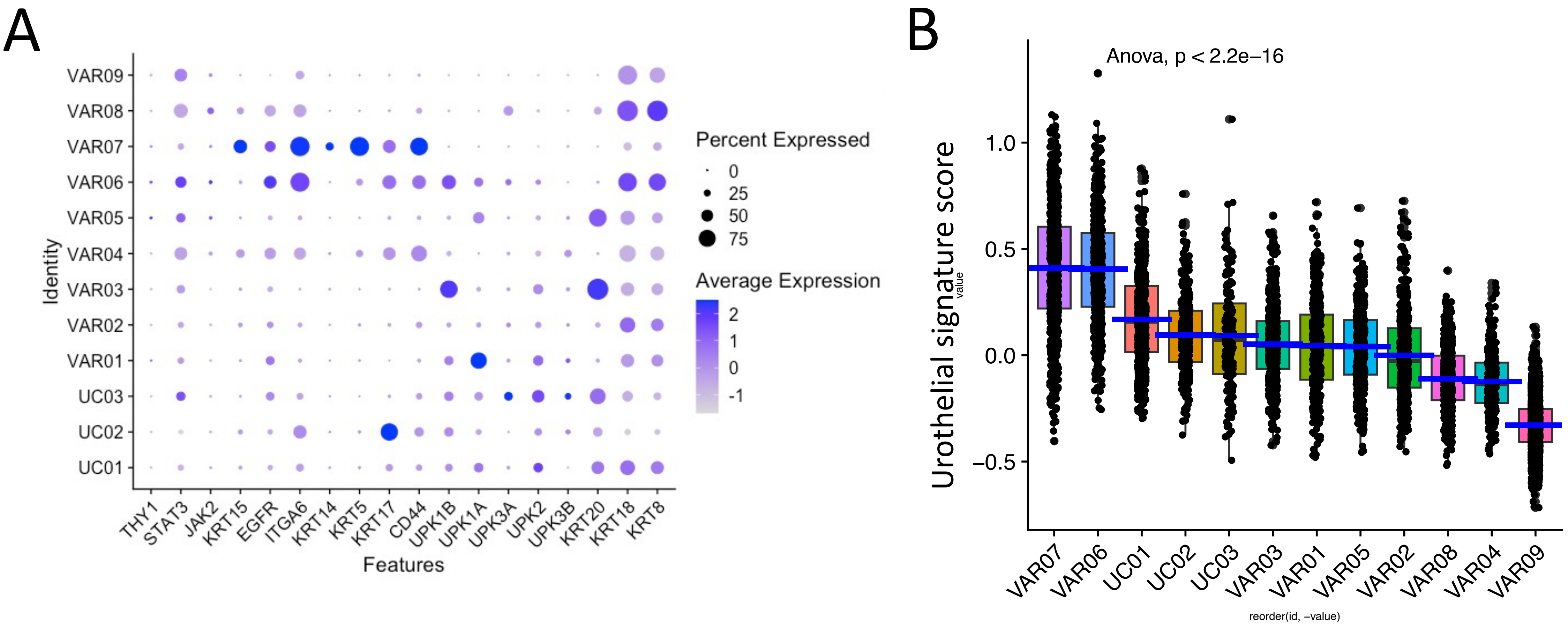

**Figure S8. Assessment of HOX gene expression among variant subclusters**

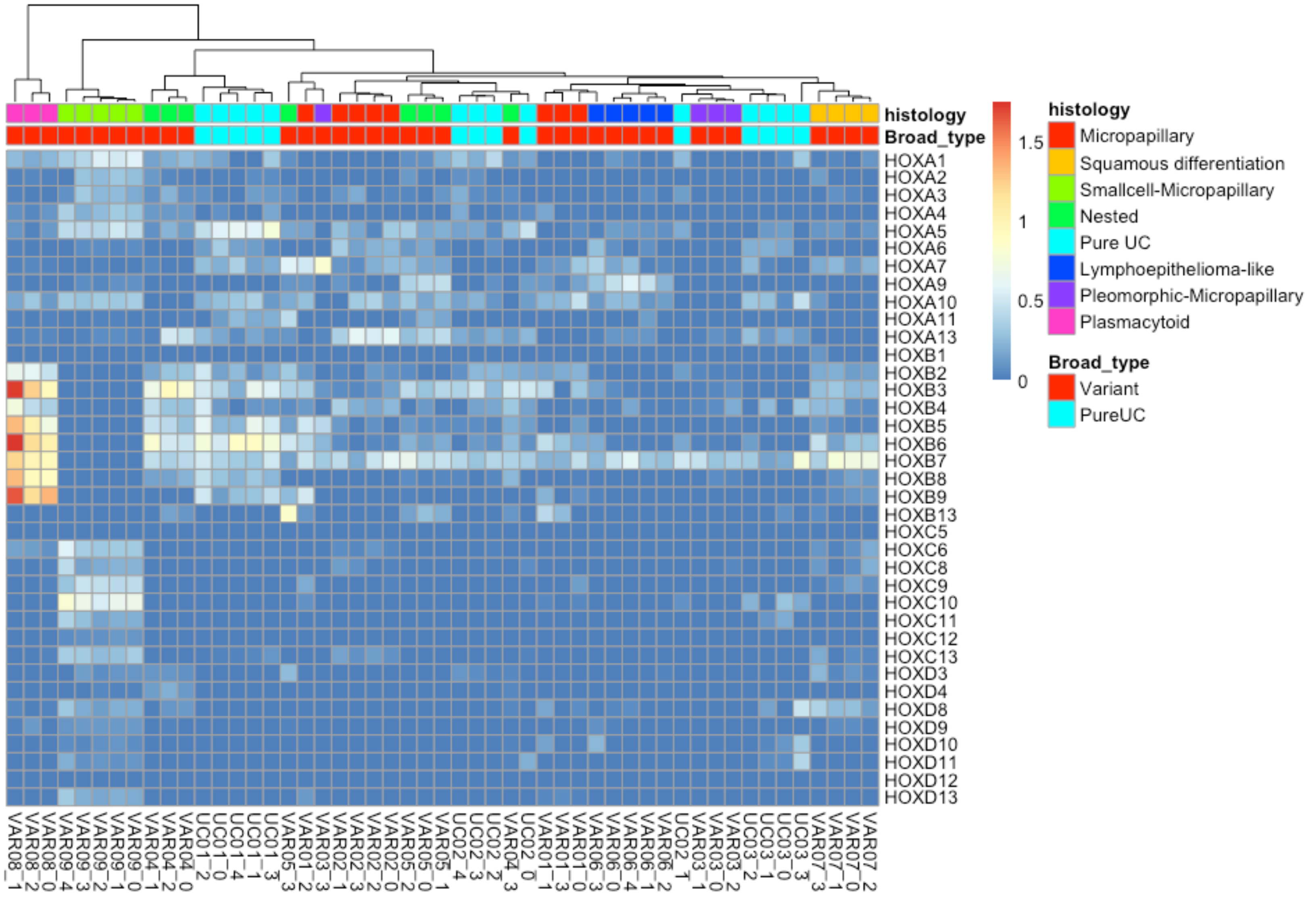

**Figure S9. Immune cell annotation and signature enrichment in tumor cells.** (A) Clustering UMAP of immune cells with top-tier annotations. (B) Heat map demonstrating Top 5 DEGs in each immune cluster. (C) Enrichment of immune cell signatures within each tumor.

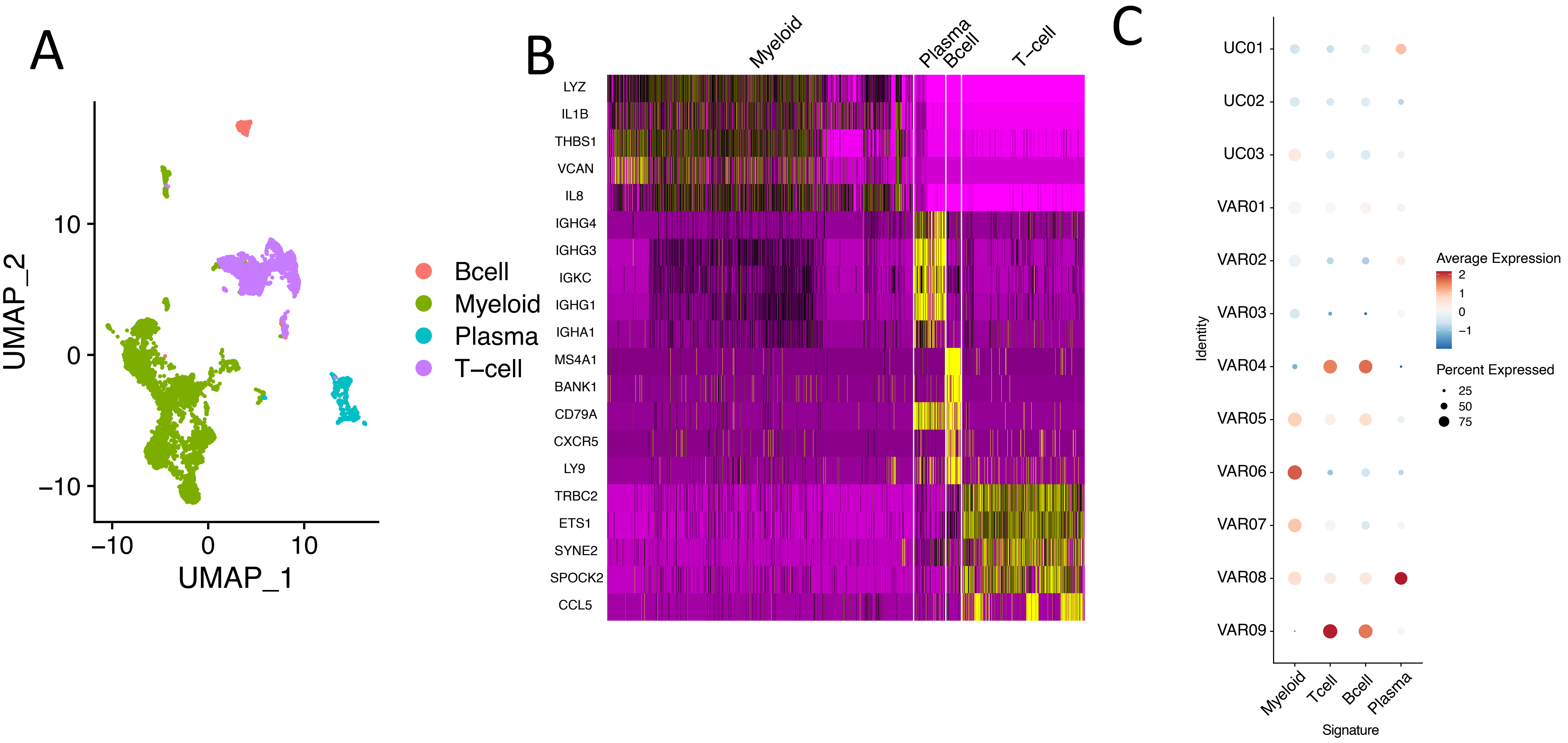

**Figure S10. Plasma cell transcriptional programs in tumor cells.** (A) Dot plot showing expression of top 100 plasma cell genes in each tumor. (B) Gene set enrichment analysis of protein secretion and unfolded protein response in VAR08 tumor cells.

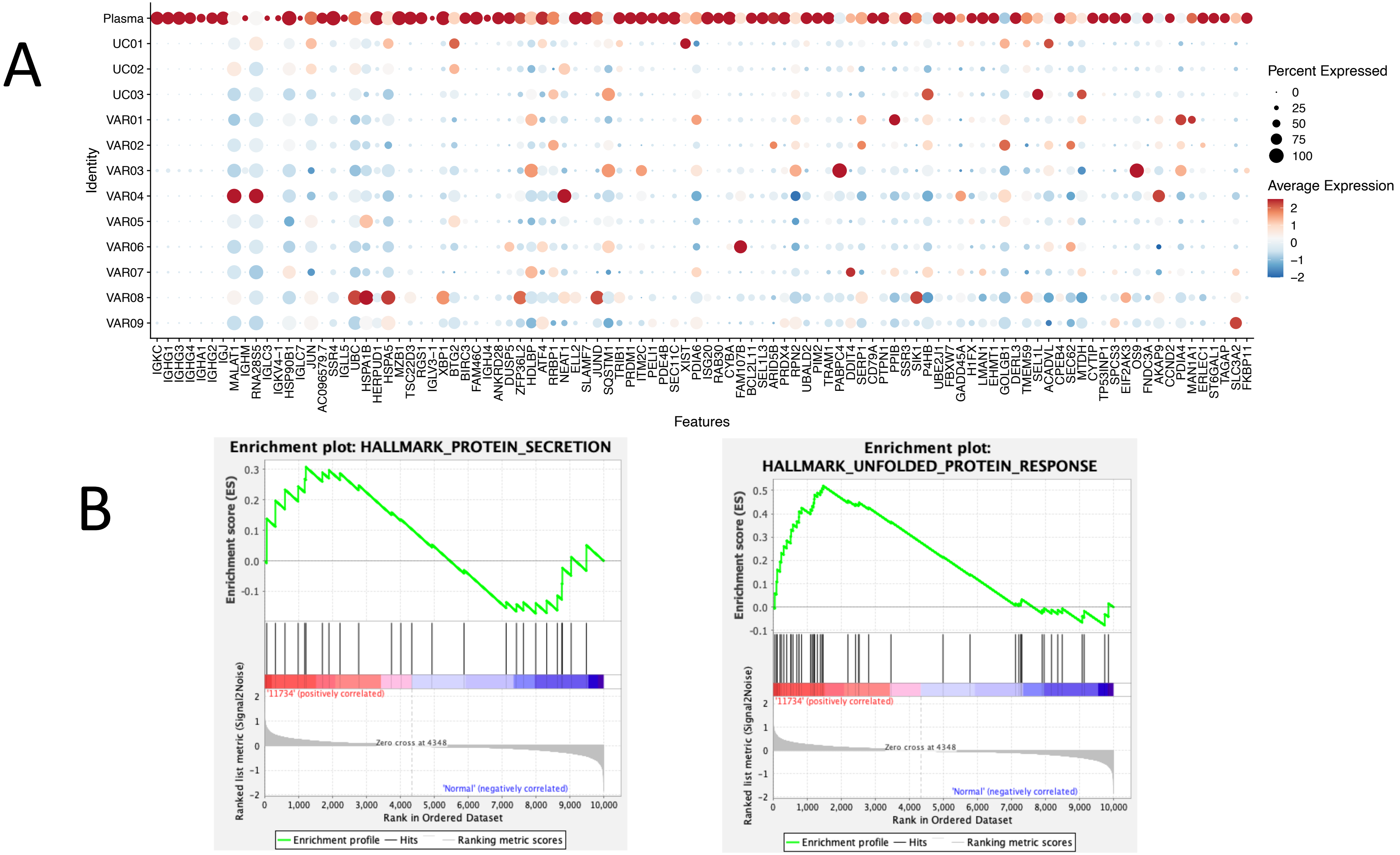

**Figure S11. Association of TM4SF1 with clinical features and luminal-basal subtypes in TCGA-BLCA.** (A) Single set gene set enrichment analysis (ssGSEA) for TM4SF1 in TCGA-BLCA. (B) TM4SF1 expression stratified by grade, stage, and lymph node status. (C) Kaplan-Meier curves showing overall survival according to TM4SF1 expression (pink = high, blue = low) in urothelial cancers, renal cancers, and pancreatic cancers.

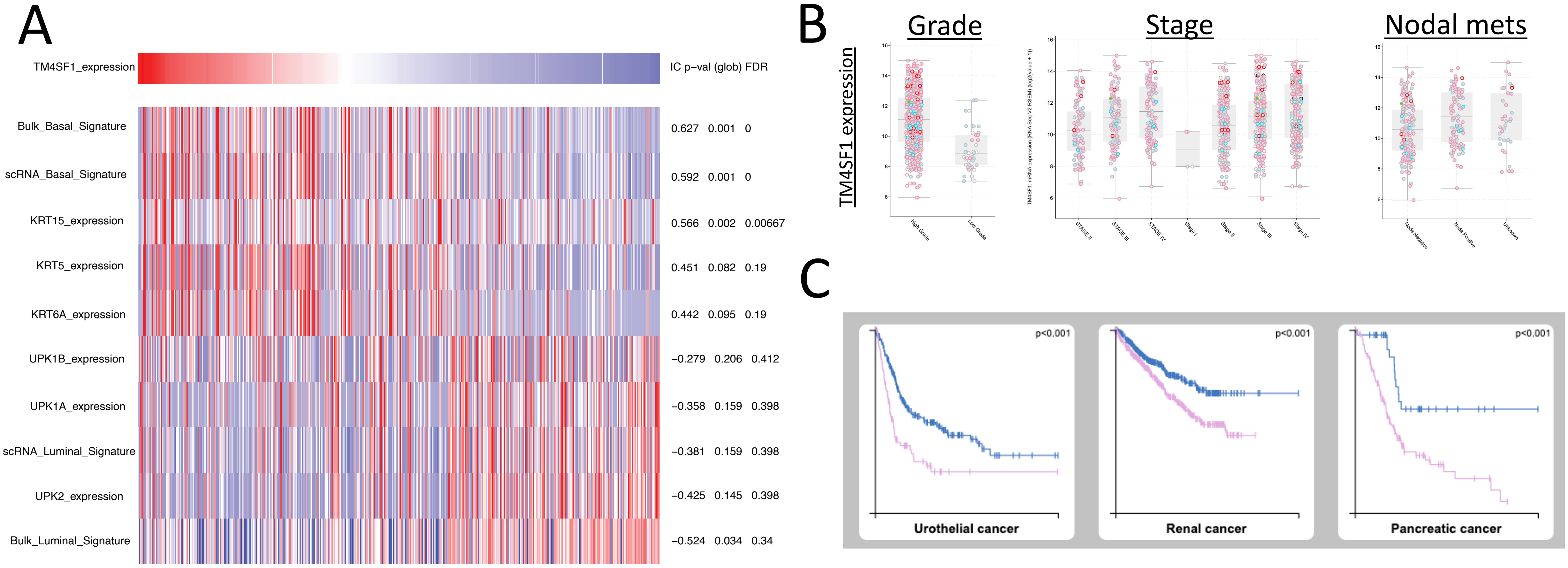

**Figure S12.** Genes correlated with TM4SF1. (A) Scatter plot showing correlation coefficient and expression of genes associated with TM4SF1. (B-C) Correlation plots between TM4SF1 and CLDN4, EZR, EMP1, and KRT19 with linear regression within (B) tumor epithelial dataset and (C) by tumor. (D) ssGSEA in TCGA-BLCA between TM4SF1 and EMP1, CLDN4, EZR, and KRT19.

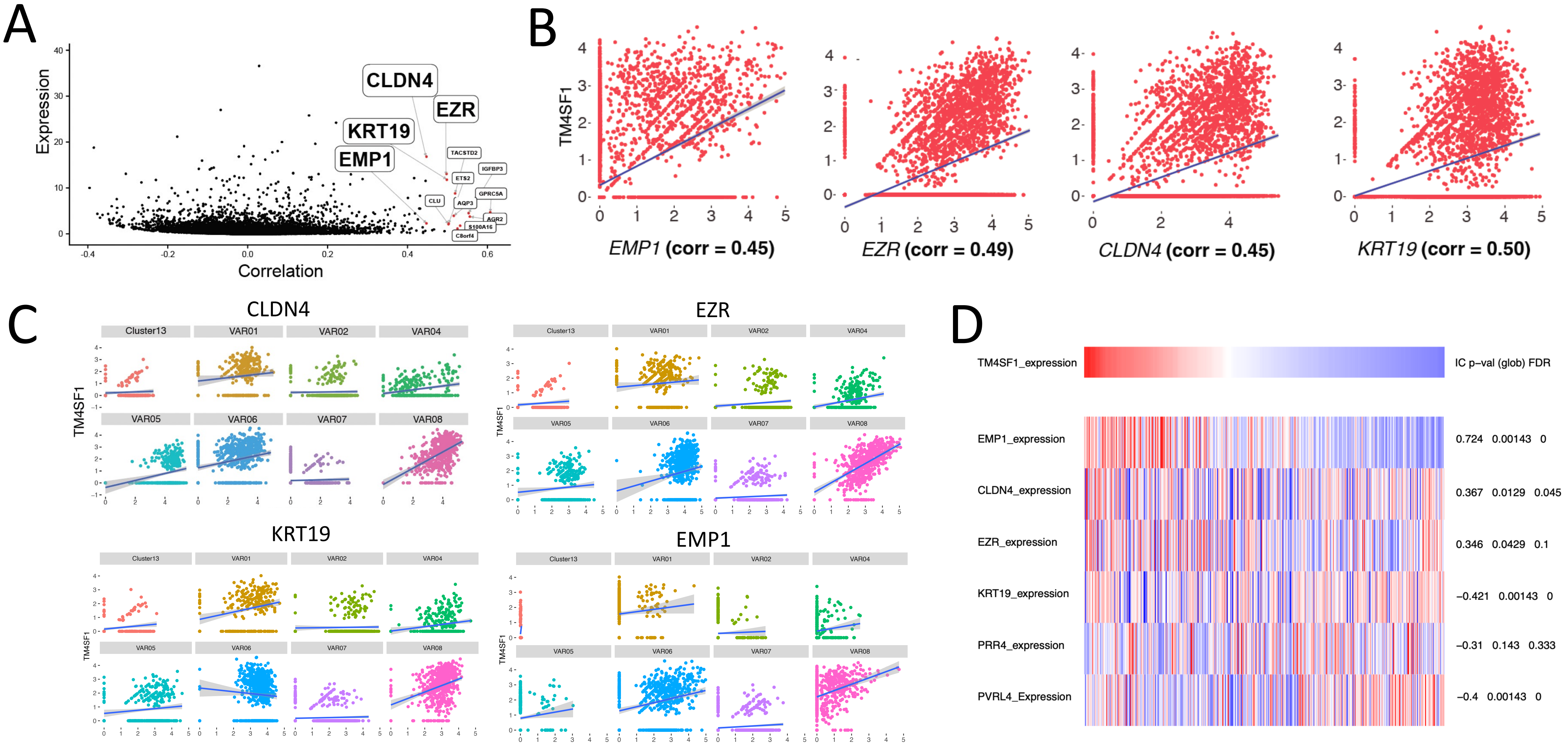

**Figure S13. Correlation between NECTIN4 and TM4SF1 in CCLE.**

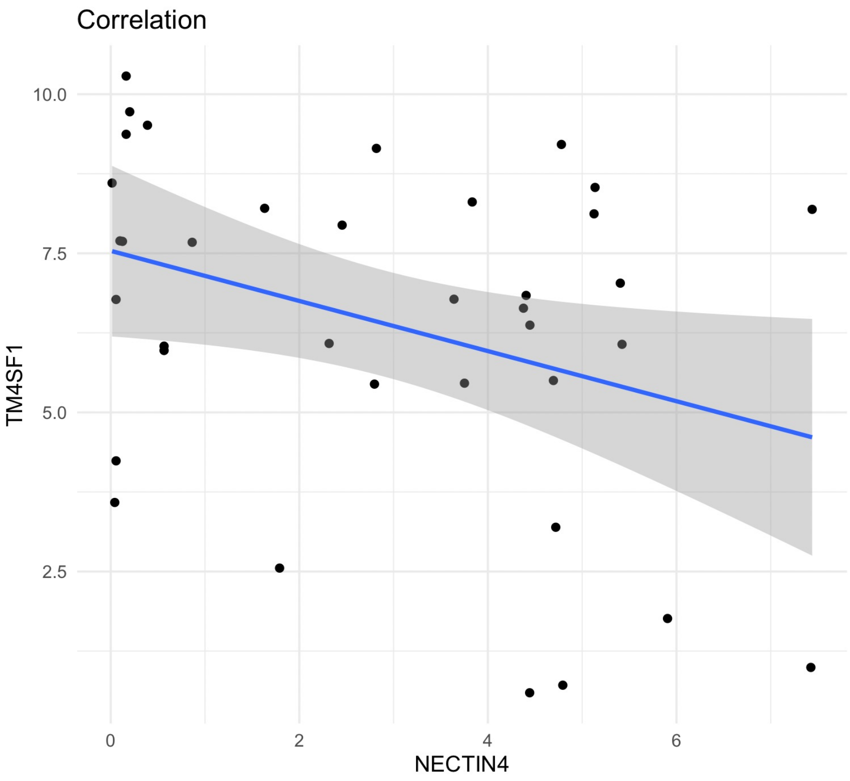


Cor = -0.35
p = 0.04

**Figure S14. CAR T cell activity against UMUC3 after knockdown of TM4SF1 using CRISPR-Cas9 .** Incucyte CAR T co-culture assay with (A) untransduced UMUC3 cells and (B) UMUC3 cells with CRISPR-Cas9-mediated knockdown of TM4SF1. A 2:1 effector:tumor cell ratio was used.

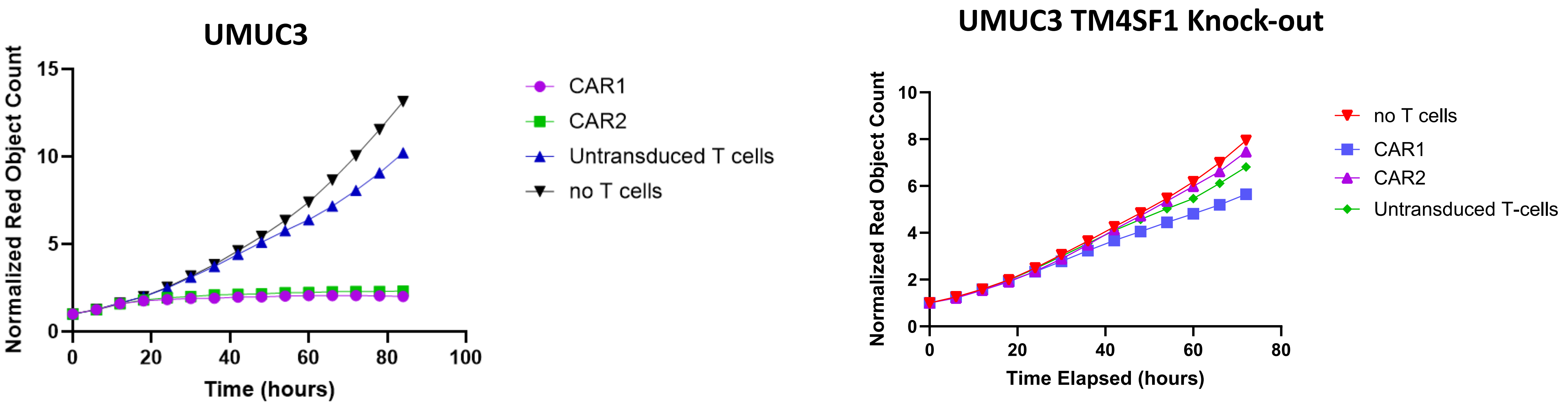

**Figure S15. Schematic for xenograft experiments**

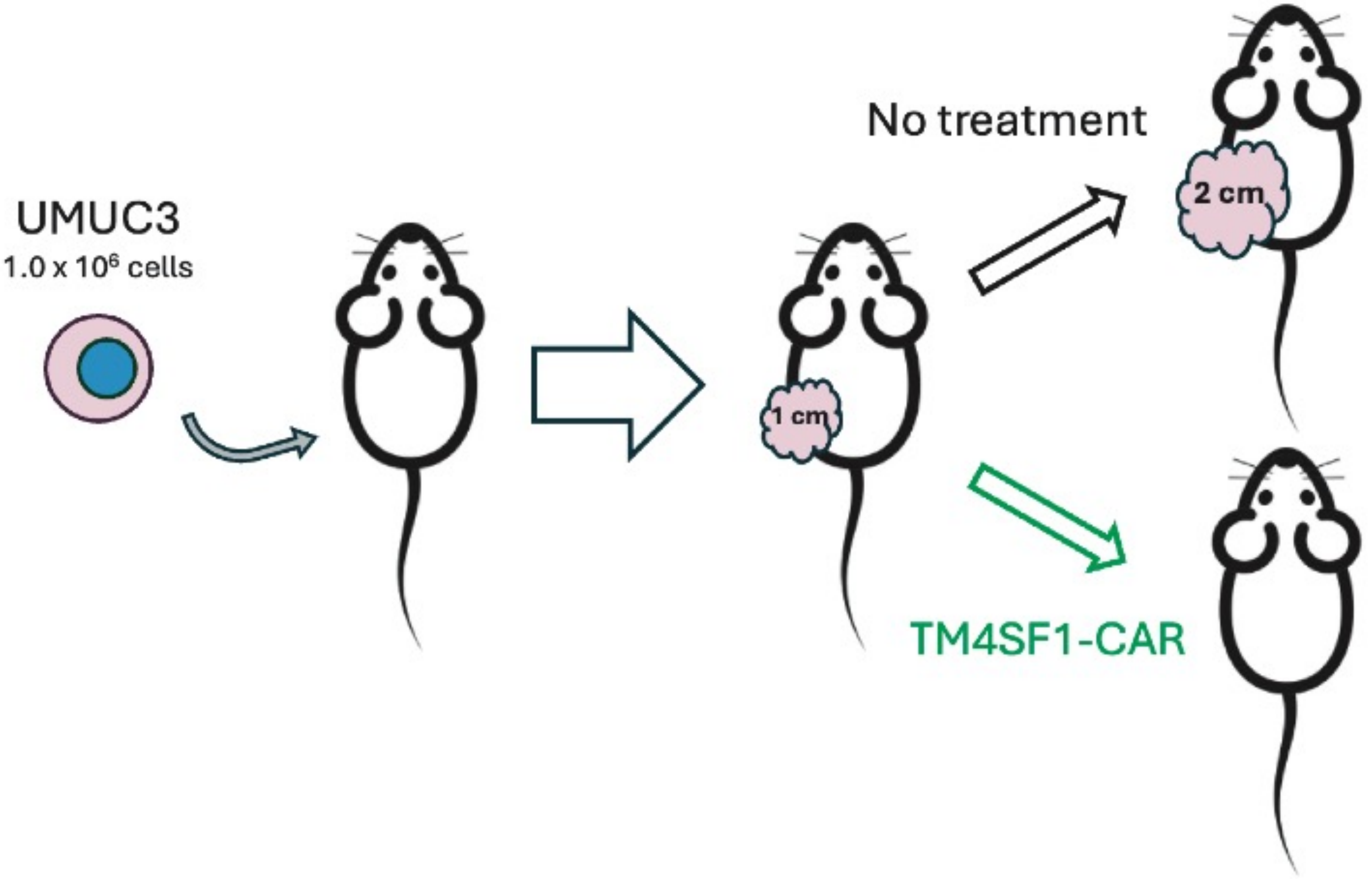

**Figure S16. Body weights of TM4SF1-CAR T treated mice.**

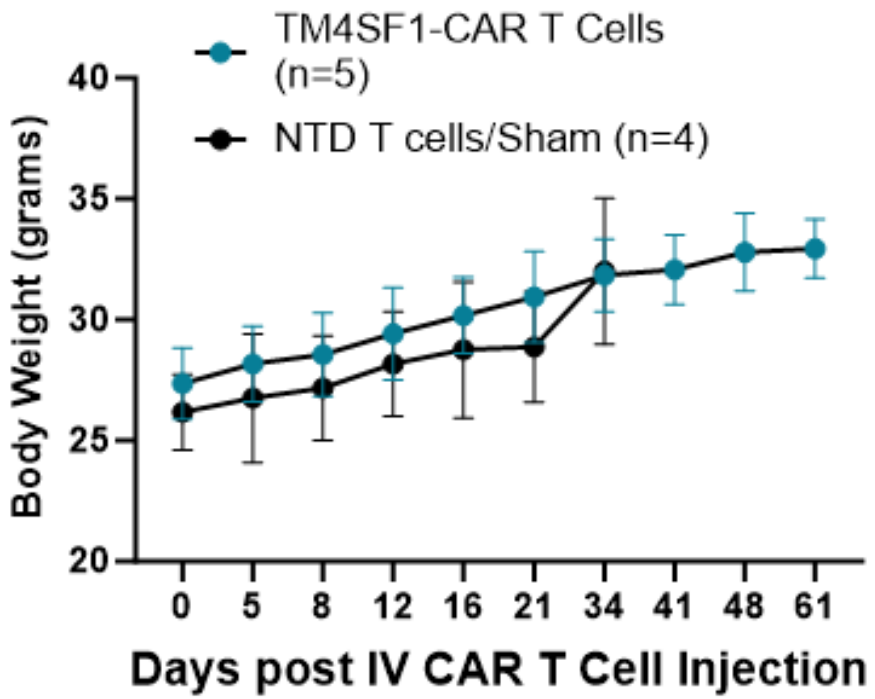
